# Improving the suppressive power of homing gene drive by co-targeting a distant-site female fertility gene

**DOI:** 10.1101/2023.12.07.570117

**Authors:** Nicky R. Faber, Xuejiao Xu, Jingheng Chen, Shibo Hou, Jie Du, Bart A. Pannebakker, Bas J. Zwaan, Joost van den Heuvel, Jackson Champer

**Author notes:** Correspondence (NRF), (JC). These authors contributed equally to this work and may list themselves as first author when referencing this manuscript (Nicky R. Faber, Xuejiao Xu).

## Abstract

Gene drive technology has the potential to address major biological challenges, including the management of disease vectors, invasive species, and agricultural pests. After releasing individuals carrying the gene drive in the target population, suppression gene drives are designed to spread at a rapid rate and carry a recessive fitness cost, thus bringing about a decline in population size or even complete suppression. Well-studied homing suppression drives have been shown to be highly efficient in *Anopheles* mosquitoes and were successful in eliminating large cage populations. However, for other organisms, including *Aedes* mosquitoes, homing gene drives are so far too inefficient to achieve complete population suppression, mainly due to lower rates of drive conversion, which is the rate at which wild type alleles are converted into drive alleles. Low drive conversion is also a major issue in vertebrates, as indicated by experiments in mice. To tackle this issue, we propose a novel gene drive design that has two targets: a homing site where the drive is located and drive conversion takes place (with rescue for an essential gene), and a distant site for providing the fitness cost for population suppression (preferably a female fertility gene, for which no rescue is provided). We modeled this design and found that the two-target system allows suppression to occur over a much wider range of drive conversion efficiency. Specifically, in the new design, the suppressive power depends mostly on total gRNA cutting efficiency instead of just drive conversion efficiency, which is advantageous because cut rates are often substantially higher than drive conversion rates. We constructed a proof of concept in *Drosophila melanogaster* and show that both components of the gene drive function successfully. However, embryo drive activity from maternally deposited Cas9 as well as fitness costs for female drive heterozygotes both remain significant challenges for two-target and standard suppression drives. Overall, our improved gene drive design eases the development of strong homing suppression gene drives for many species where drive conversion is less efficient.

## Introduction

Gene drive technology has the potential to address major biological challenges, including the management of disease vectors, invasive species, and agricultural pests(1–5). A gene drive is a genetic element that skews its own inheritance ratio to over 50% in the offspring of heterozygote parents(6). Synthetic gene drives can be designed for population modification, for example to immunise a population of mosquitoes against malaria parasites(7), or for population suppression, for example to eliminate a population of mosquitoes(8) or invasive pests(9, 10). Released in the target population, suppression gene drives are designed to spread at a rapid rate and carry a recessive fitness cost, thus causing a decline in population size or even complete elimination(8, 11, 12). Suppression gene drives have benefits over conventional methods of control because they are species-specific (and thus more ecologically friendly), as well as potentially more efficient and more humane in the case of vertebrates, though there are challenges regarding localisation and containment for some drive types(13).

There are many different types of gene drive and what distinguishes them the most is how they handle the trade-off between efficiency of spread and confinement(2, 5, 6). The most efficient and well-studied type of suppression drive is the CRISPR-Cas9-based homing drive. In drive heterozygotes, the gene drive copies itself to the homologous chromosome in the germline through a process called “homing” or “drive conversion”. The gene drive incurs a recessive fitness cost by being located inside a haplosufficient female fertility gene, so drive homozygous females are sterile. Ideally, the female infertility is completely recessive, so there are no fitness costs for drive heterozygotes. As the frequency of the gene drive increases, more sterile female offspring are created, thus imposing a genetic load on the population(14). If the frequency of sterile females in the population is sufficiently high, the population size will decline(2, 9).

Although homing gene drives are very promising in theory, *in vivo* tests in several organisms have revealed practical challenges, so complete population suppression is not yet attainable in most species (see Figure 1)(9, 13, 15–20). Homing suppression gene drives face two major challenges, the first being formation of functional resistance alleles(16, 20– 22). A typical gene drive is active in the germline, and the primary way in which Cas9 cuts will be repaired is through homology-directed repair(23). In this pathway, the cell uses the drive chromosome as a template, copying its sequence to the other chromosome. Sometimes, particularly if the cut is made outside of the time window in which this repair pathway is preferred, repair can occur via end-joining instead. This repair pathway is error-prone and often leads to small insertions and/or deletions. Because of these mutations, the gene drive guide RNA (gRNA) usually cannot recognise the new sequence, making it a resistance allele that can no longer be converted to a drive allele. Resistance alleles that preserve the function of the target gene, or “r1” alleles, are problematic as there is very strong selection for this allele which will rescue the population from the suppression drive(16). Nonfunctional resistance alleles (“r2” alleles), on the other hand, will not be able to rescue the population because of their deleterious nature(20). Although r2 alleles can reduce drive efficiency, the less common r1 alleles are the bigger problem because they will outcompete the gene drive in the population. There are several strategies to avoid the formation of r1 alleles in suppression drives, which include targeting an extremely conserved locus(8), multiplexing gRNAs(19, 24, 25), and using improved promoters(26–28).

**Figure 1.**
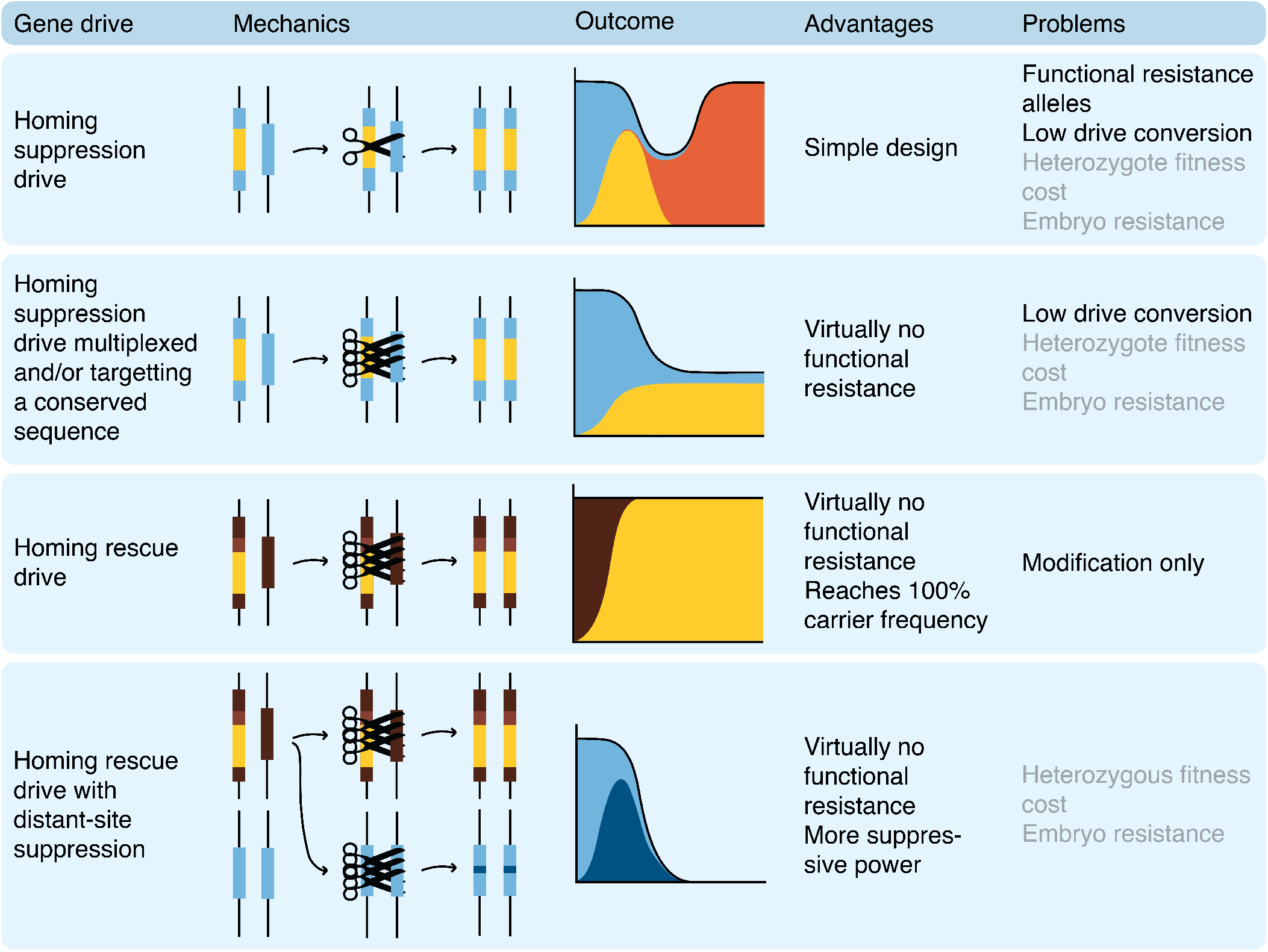
Overview of several homing gene drive designs, their advantages, and major challenges they face. In the outcomes graphs, light blue = female fertility gene, yellow = gene drive, orange = functional resistance allele, dark brown = essential gene, dark blue = non-functional resistance allele (only the female fertility target site is displayed for the distant-site suppression system). Additionally in the schematics, light brown = essential gene rescue. Primary problems for drive designs are in black, and secondary problems in grey.

Besides r1 alleles, a second challenge for suppression gene drives is low drive conversion, which is the rate at which wild type alleles are converted into drive alleles(17, 25). In *Anopheles* mosquitoes, drive conversion has been shown to reach 95-99%(8, 29). In *Aedes* mosquitoes and *Drosophila melanogaster* on the other hand, the conversion rate was usually significantly lower, around 50 to 70% for most constructs(20, 25, 30–32). Additionally, it appears that multiplexing, which is a way to avoid r1 alleles, may further reduce conversion rates beyond 2-4 gRNAs(19, 25). This low efficiency results in partial population suppression instead of population elimination. The amount by which the population size is reduced compared to the expected population size in the absence of the gene drive is the genetic load of the gene drive when its reaches its equilibrium frequency in the population(14). In an infinitely large population, the genetic load needed for elimination is one minus the inverse of the low-density growth rate (the reproductive advantage individuals experience in the absence of competition), but in smaller or less fecund populations, genetic drift and stochasticity can somewhat ease this requirement. Drive conversion efficiency seems to be partially based on the species for the most commonly used promoters, though improvements might be made by regulating the expression of the gene drive to more precisely coincide with the window for homology-directed repair(17, 26, 28, 33, 34), or perhaps by more strongly localising the gene drive mRNA to the nucleus(35).

To tackle this issue, we propose a novel gene drive design that has two targets: a homing site where drive conversion takes place, and a distant cutting site (where the drive is not present) for providing the fitness cost for population suppression (see Figure 1). For the homing site, we will harness a modification drive that is located in an essential gene (thereby disrupting it) while also providing a rescue for this gene (see Figure 1)(36). A previous study has already demonstrated the practical feasibility of using a homing gene drive while also targeting another gene for population modification(37), and another recent study has shown that modifying a natural gene drive to also target a distant female fertility gene is feasible for population suppression(10). Thus, we model the two-target design, comparing it to standard homing suppression gene drives, and find that this two-target system allows suppression to occur over a much wider range of drive conversion efficiency. Most notably, the suppressive power now depends usually on cutting efficiency at the distant target instead of drive conversion, which is advantageous because the total cutting rate is easier to increase and has often been substantially higher than drive conversion rate(17, 18, 20, 32). We constructed a proof of concept of this drive in *Drosophila melanogaster* and show that both components function successfully. The drive conversion rate was within the range generally observed in *D. melanogaster*, and the cut rate at the distant-site target was very high. However, high fitness costs still thwarted the drive’s success in cage populations, a factor that could potentially be problematic in any suppression drive based on targeting essential genes. Nevertheless, our improved gene drive design enables development of strong homing suppression gene drives for a wide array of species where drive conversion is less efficient.

## Methods

### A. Modelling

We use a stochastic, individual-based model with non-overlapping generations in a randomly mating population of fixed carrying capacity. We use the population genetics modelling software SLiM version 4.0.1 (38) in combination with R version 4.1.0 (39). Our code can be found on GitHub at https://github.com/NickyFaber/Two-target_drive.

#### A.1. Drive types

We have modelled 10 different types of homing suppression drives. For ease of visualisation, we have chosen to show the three most representative drives in the main results. A list of the rest of the drives and their modelling results are in the Supplementary Materials.

1. Drive with female fertility target
2. Haplolethal rescue drive with distant-site female fertility target
3. Haplosufficient rescue drive with distant-site female fertility target

For a detailed explanation of the mechanics and phenotypes of these three drives, see the results. To model each drive, we include two loci in SLiM: the homing site and an optional distant-site. At the homing site, there are four potential alleles: wild type, drive, r1 (functional resistance), and r2 (nonfunctional resistance). At the distant site, we only model wild-type and r2 alleles (full freedom to choose any set of gRNAs is assumed to reduce functional r1 resistance to negligible levels, see results). The structure and steps of the model are described below.

#### A.2. Model structure

We have used previous work by Champer et al. (2020) as a starting point for our modelling(19). That study modelled complex drive activity in the germline, including gRNA multiplexing, timing of drive activity, and gRNA saturation. Since our objective is to compare various gene drive designs focusing on genetic load instead of on resistance alleles, we have removed some of those complexities for this study. We model the population and introduction of the gene drives as follows:

- Generation 1: Population initialization
- Generation 1-10: Population equilibration without gene drive
- Generation 11: Introduction of heterozygous gene drive individuals
- Generation 111 or 161: End of the model. As default, we run the model for 111 generations (100 generations with gene drive). When we calculate the genetic load, we run it for an additional 50 generations to increase the number of generations in which we can determine the equilibrium genetic load.

In each generation, the model executes the following steps in order (see Figure 2):

**Figure 2.**
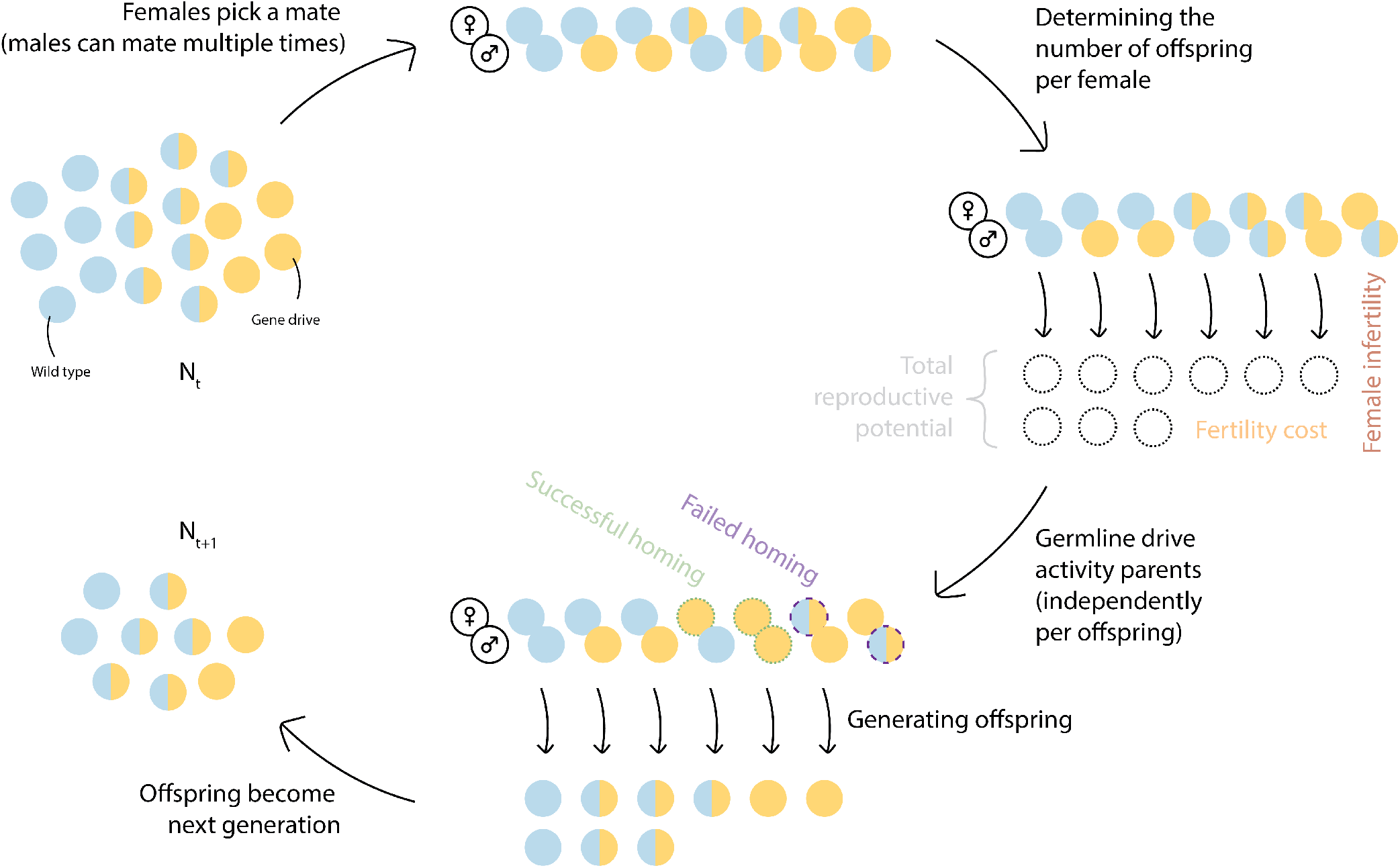
Model overview, showing a simplified sequence of steps the model goes through every generation. Each circle represents an individual, and its color shows the genotype. We show a simple homing gene drive with only wild type (blue) and drive (yellow) alleles. The two-target drives will show different dynamics dependent on their design (see figure 3). After each female picks a male for mating, we calculate the reproductive potential of these females. Then, based on drive genotype, females can be either completely fertile, or completely sterile (drive homozygotes), or partially fertile (drive heterozygotes). Offspring is generated, and for each individual offspring, Mendelian inheritance is modified by drive activity in the parental germline. This means that drive conversion could be successful for one offspring of a single drive parent, but not another. Additional drive activity from maternally deposited Cas9 is also modeled, but not shown in this figure. Finally, these offspring become the next generation of adults.

##### 1) Reproduction

In our model, the number of offspring that is generated by each female is a product of the fertility status of the female, her ability to find a mate, the carrying capacity, and the fitness of the female.

- Fertility status check. A female could be infertile due to the drive mechanism, both at the homing site and the distant site. These loci are both checked for the presence of nonfunctional alleles, that is, either a drive allele or an r2 allele. If the female has two of these in any combination at at least one locus, she does not generate any offspring.
- Selecting a mate. A female randomly selects a male from the population. If there are no males available, she does not generate any offspring.
- Generating offspring. For each female, the number of offspring she generates in that generation (*i*) is based on a binomial distribution:

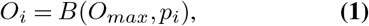

where *i* is the current generation, *O*_*max*_ is the maximum amount of offspring per female, and *p* is the average fraction of this maximum amount of offspring that will be generated. This fraction is normally defined as 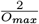 (each female must generate two off-spring to maintain the population at carrying capacity). However, our model includes two further influences. First, *p* can be increased when the population is below carrying capacity because offspring will have more resources to survive, or vice versa, which we call the carrying capacity factor (*CCF*). Second, *p* can be decreased in female drive carriers due to somatic expression of the drive reducing fertility (because some wildtype female fertility alleles are disrupted in somatic cells where they are needed for fertility), which we call the somatic expression fertility factor (*SEFF*). Thus, *p* is defined as:

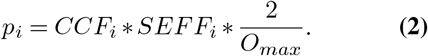

The carrying capacity factor is defined so that at very low population densities, it is close to the maximum growth rate (*r*), at carrying capacity (*K*), it is close to 1, and above carrying capacity, it is between 0 and 1. This leads to a logistic growth curve:

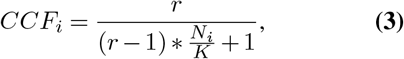

where *N* is the number of adults in the population. Fertility scaling is done for females with at least one drive allele. Somatic expression of the drive can impact female fertility by prematurely disrupting wildtype fertility gene alleles. One of our modelled drives targets a female fertility gene at the homing site without rescue and the other two at the distant site. We only model somatic expression fitness costs for the drive sites without rescue. The total fertility cost is calculated per female as follows:

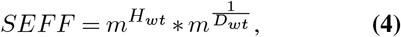

where *m* is the somatic expression fertility cost multiplier, from 0 (complete sterility) to 1 (no fertility cost), *H*_*wt*_ is the number of wild type alleles at the homing female fertility site (which can only be 0 or 1, since the other allele must be the drive), and *D*_*wt*_ is the number of wild-type alleles at the distant female fertility target. At the distant locus, individuals with two wild-type alleles will be less impacted by the fertility cost, since it is less likely that both copies of the fertility gene are disrupted due to somatic expression.

##### 2) Gene drive activity

After all the offspring is generated, each offspring’s genotype is modified based on the drive activity in the parents’ germline. Drive activity in the parental germlines is modelled for each offspring independently. Cutting, homing, and the creation of resistance alleles is stochastic.

- Germline gene drive activity. For both parents of each offspring, the presence of the drive in their genome is checked. If at least one copy of the drive is present, the gene drive is active in the germline. First, the cut rate is a parameter in our model, but it can be impacted by gRNA saturation as follows:

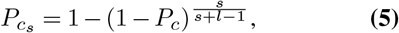

where 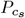 is the cut rate adjusted for gRNA saturation, *l* is the number of loci, either 1 or 2 depending on whether the drive targets a distant site in addition to the homing site (we assume equal gRNA multiplexing at both sites), *P*_*c*_ is the global cut rate, and *s* is the Cas9 saturation factor, which can range from 1 (which means 1 gRNA is enough to saturate all Cas9 proteins, so the cut rate for each gRNA rapidly declines as more gRNAs are added) to infinite (which means no amount of gRNAs is enough to saturate the Cas9 proteins, so the cut rate at each gRNA remains the same regardless of the number of gRNAs). If a randomly generated number between 0 and 1 is higher than this cut rate, cutting occurs. We do this separately for the distant site alleles as well, if present. At the distant site, cutting always results in an r2 allele. At the drive locus however, homing can occur if there was successful cutting based on the homing success rate (*P*_*h*_). The homing success rate is defined at the beginning of the model based on the conversion rate (*P*_*conv*_), which is a parameter:

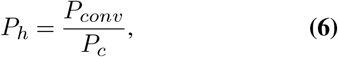

where *P*_*conv*_ can only be as high as *P*_*c*_. If homing is successful, again based on *P*_*h*_, the allele is converted to a drive allele. If there was cutting, but no homing, the locus is converted into an r1 allele with a probability equal to the r1 formation rate. Otherwise, it will become an r2 allele.
- Embryo gene drive activity. In the early embryo, we model maternal deposition of drive Cas9 and gRNAs. Potential cuts that occur here always result in a resistance allele (which can be r1 or r2 as above). The cut rate 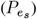 is calculated the same as in Formula 5, except the exponent is additionally multiplied by a maternal deposition factor (*d*) that accounts for either a drive heterozygous or homozygous mother:

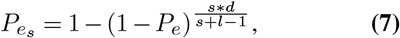

where *P*_*e*_ is the embryo cut rate, which is a parameter in our model. The maternal deposition factor is based on experimental data showing that the cut rate is higher in embryos with a drive/wild-type heterozygous mother than expected due to drive conversion, so *d* is 1.83 and 2 for drive/wild-type heterozygotes and drive homozygotes, respectively. It would remain 1 for fertile drive/resistance allele heterozygotes(19).
- Offspring viability. After drive activity in the parental germline, inheritance, and embryo activity, we check if the offspring’s resulting genotype is still viable. All drives that target either a haplolethal gene (where any nonfunctional resistance allele makes individuals nonviable) or haplosufficient gene (where only nonfunctional resistance allele homozygotes are nonviable) could result in nonviable offspring. These offspring are removed from the population.

##### 3) Mortality

We model discrete, non-overlapping generations, so we remove the entire parental generation after offspring have been generated.

##### 4) Calculating genetic load

Because our model is stochastic, populations can be suppressed if the genetic load is close to, but lower than the deterministic requirement, which is below 1 and depends on the maximum growth rate of the population. Therefore, in order to calculate genetic load with precision for drives with high genetic loads, we run a module in the model in which we artificially raise the number of offspring a female produces by multiplying it with a certain bonus factor(19). These are later corrected for when we calculate the genetic load. With this approach, the population is not eliminated unless the genetic load is practically 1, allowing for precise measurements of high genetic load for several generations when the drive is at its equilibrium frequency. This bonus factor (*BF*) is calculated as follows:

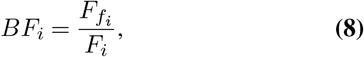

where *F* is the number of females, and *F*_*f*_ is the number of fertile females.

Then, in the next generation, the number of offspring calculated in Formula 1 is increased as follows:

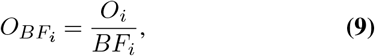

rounded to a whole number. Additionally, in genetic load simulations, gene drive carriers are introduced at 0.5 frequency, and the model is run for 150 generations after introducing the gene drive. The mean genetic load is calculated as the mean genetic load over the last 10 generations. The lowdensity growth rate and the maximum number of offspring are set a factor 10 higher than default (so 100 and 500 instead of 10 and 50, respectively).

##### 5) Tracking outcomes of interest

Each generation, we calculate population size, genetic load, and genotype frequencies at both the homing site and the distant site. We calculate the genetic load (*GL*) based on the observed and expected population size in the next generation (*N*_*i*+1_ and 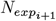, respectively):

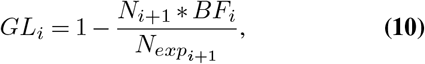

where *BF* is the above mentioned bonus factor we apply.

*N*_*exp*_ is based on the number of females (*F*) and the carrying capacity factor *CFF* defined in Equation 3:

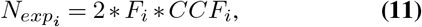

where *CFF* is the same carrying capacity factor defined in Equation 3, and again multiplying by 2 because each female must generate two offspring to maintain the population at carrying capacity.

### B. Experimental work

#### B.1. Plasmid construction

The starting plasmids TTTgR-NAt and TTTgRNAtRNAi were used for building gRNA helper plasmids for knock-in(36). The gRNA cassette used in the donor plasmid was obtained from HSDygU4(25). A two-step assembly process was done to generate donor plasmids (SI Appendix, Methods). Q5 High-Fidelity DNA Polymerase used for PCR and enzymes for digestion were purchased from New England Biolabs. PCR and restriction digestion products were purified with Zymo Research Gel DNA Recovery Kit, and plasmids were assembled by using HiFi DNA Assembly Cloning Kit and subsequently transformed into DH5α competent cells from TIANGEN. ZymoPure Midiprep Kit (Zymo Research) was used to prepare donor constructs for embryo injection. Oligo synthesis and sanger sequencing was done by BGI Genomics. All the primers, plasmids, and construction procedures used in this study can be found in can be found in SI Appendix, Methods. Plasmid maps are available on GitHub (https://github.com/NickyFaber/HaploLethalFertilityDrive) in ApE format(40).

#### B.2. Generation of transgenic lines

Fly injection was completed by UniHuaii. The donor plasmid HSDrgU2U4v2 (518 ng/ul) was injected into AHDr352v2 flies(36) along with TT-TrgU2t (150 ng/ul), which provided gRNAs for transformation, and Cas9-expressing helper plasmid TTChsp70c9 (450 ng/ul) to generate the drive line. The other donor plasmid SNc9NR (506 ng/ul) was injected into w1118 flies along with helper plasmids BHDabg1 (100 ng/ul) and TTChsp70c9 (459 ng/ul) to construct a *nanos*-Cas9 line. Surviving G0 flies were crossed to w1118 flies, and G1 adults were screened for transgenic inserts based on the presence of green or red fluorescence in the eyes. Flies were reared in an incubator at 25°C following a 14/10-h day/night cycle.

For phenotyping, flies were first anesthetized with CO_2_ and then screened for fluorescence using the NIGHTSEA system (SFA-GR). The homozygosity of flies was scored by the fluorescence intensity and confirmed by sequencing.

#### B.3. Drive conversion

Drive (gRNA-expressing line) males were crossed to Cas9 females to generate heterozygous offspring, which was subsequently out-crossed to *w1118*. The drive inheritance and sex of offspring were recorded. To confirm whether the distant target site in yellow-g was disrupted, individuals containing both drive and Cas9 alleles were randomly collected for genomic DNA extraction and genotyping. A fragment covering yellow-g target sites was amplified with primers 52_YGLeft_S_F3 and 54_YGRight_S_R6 (see GitHub DNA files).

#### B.4. Small cage study

Drive heterozygous males with red fluorescence from the Cas9 allele were crossed to homozygous Cas9 females for several generations to produce a line that was heterozygous for the drive and homozygous for Cas9 (D/+; Cas9/Cas9). Two experimental groups were set up with different initial release frequencies. In the higher release frequency group, four drive females were crossed to four drive males, while in the medium release frequency group, four drive females were crossed with four drive males, and one Cas9 virgin female was crossed to Cas9 males. Thus, the high drive frequency release should theoretically be 0.5 (1.0 carrier frequency), and the medium drive frequency release 0.4 (0.8 carrier frequency). These adults were allowed to mate in vials for one day before moving females into a separate bottle for oviposition. Females were allowed to lay eggs (which represented “generation 0”) for three days and were then removed from bottles. When most pupae eclosed to adults, they were moved to a new bottle for a one-day oviposition before being removed and phenotyped. Hereafter, only one-day oviposition was conducted in each generation. The adults of each generation were scored for eye fluorescence phenotype and sex.

#### B.5. Fecundity and fertility test

To minimize batch effects caused by food quality or population density, flies with different genotypes used for this test were generated from the same parental cross and reared in the same bottle. First, males that were heterozygous for the drive and homozygous for Cas9 were crossed to Cas9 homozygous females, generating offspring with different genotypes. Next, these offspring were individually crossed to Cas9 homozygous males or females and allowed to lay eggs for three days. Adults in the same vial were moved to a new vial each day, and the number of eggs was counted in each vial. Offspring were allowed to hatch, and the egg-to-adult survival rate and adult phenotypes of these offspring were scored. Female drive offspring were randomly collected and crossed to Cas9 males, after which the sterile females were genotyped for the *yellow-g* distant site.

## Results

### A. Modelling the two-target gene drive performance

In our main results, we show the modelling results of three gene drives: a standard female fertility homing suppression drive and two two-target drives that target a female fertility gene at a distant site for population suppression. These two-target drives are located in and provide rescue for a haplolethal or a haplosufficient gene. Drive conversion occurs normally in these, but they also cut and disrupt a distant-site female fertility target without rescue. Figure 3 shows the three drives’ loci and dynamics, and also the alleles and phenotypes that can result in gametes from drive activity.

**Figure 3.**
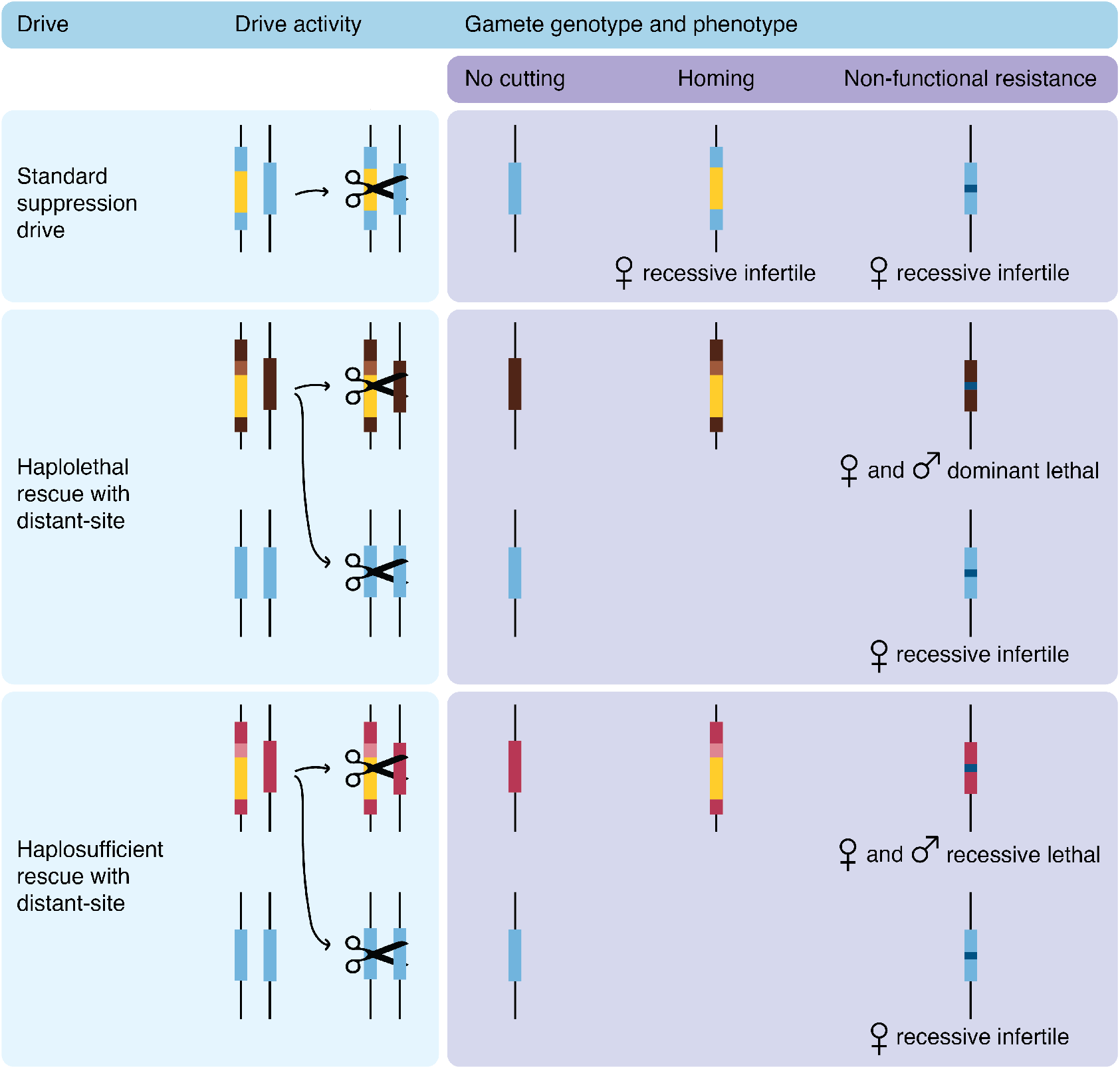
Overview of our three modelled drives, their activity, and the resulting gamete genotypes and phenotypes. We do not show functional resistance alleles, but their phenotypes would be the same as wild-type. Light blue = female fertility gene, yellow = gene drive, dark blue = non-functional resistance allele, dark brown = haplolethal essential gene, light brown = haplolethal essential gene rescue, dark pink = haplosufficient essential gene, light pink = haplosufficient essential gene rescue.

#### A. 1. Cut and conversion rate

Because the drive conversion rate is one of the most important parameters to determine the success of a suppression drive, we start by varying it, together with the total cut rate (referring to germline cutting)(12). Any wild-type alleles that are cut but not converted to drive alleles are converted to nonfunctional resistance alleles (Figure 4A). At the distant female fertility gene target, there is no drive allele present, so the total cut rate can only result in nonfunctional resistance alleles. Besides these, all other parameters are fixed at optimum values (no embryo resistance, no fitness costs, no functional resistance, and no effect of Cas9 saturation). Figure 4B shows population size over time after a gene drive introduction.

**Figure 4.**
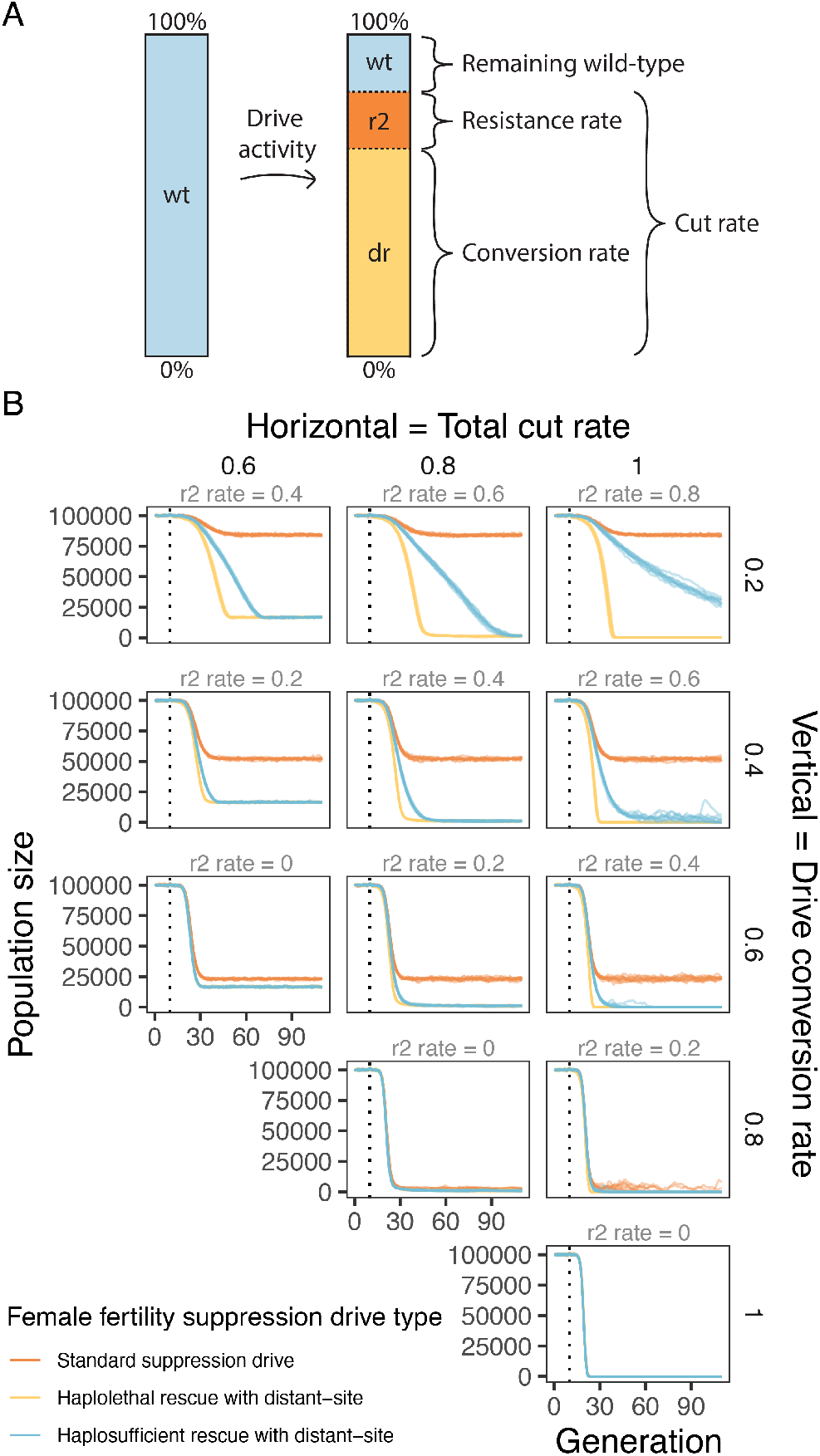
Example population trajectories under varying cut and conversion rates. **A)** Schematic of drive activity rates in the germline. **B)** Population suppression with varying total cut and drive conversion rates. The gene drive is introduced after generation 10 (the dotted line). The introduction frequency of gene drive heterozygotes is 0.01, and the total population size is 100,000. Note that both drive conversion rate and total cut rate are absolute rates, so the drive conversion rate can never be higher than the total cut rate. For each combination of parameters, we ran 10 model repetitions for each drive that are each shown as a translucent line.

At 100% cutting and drive conversion, all three drives work equally well, rapidly eliminating the population. As the conversion rate decreases to 0.8, we still observe drive success, but stochasticity now plays an important role in achieving population elimination for the standard suppression drive. The haplosufficient rescue drive with distant-site suppression experiences such fluctuations as the drive conversion is further reduced, but it still achieves rapid success when drive conversion is 0.6, at which point the standard suppression drive fails. Only the haplolethal rescue drive with distant-site suppression is still able to reliably achieve population elimination.

These major differences between drives can be explained by the standard suppression drive relying on drive conversion to both spread and suppress the population, whereas both two-target drives use two separate loci for this (Figure 3). The haplolethal distant-site drive can make r2 alleles at the drive locus that are immediately removed due to the haplolethal nature of the gene (Figure S1A). Therefore, this gene drive spreads the most efficiently. The haplosufficient rescue drive and the standard suppression drive also form deleterious r2 alleles, but these can remain in the population due to slower removal, which impairs drive spread because drive conversion will not occur in drive/r2 heterozygotes. Additionally, in the standard suppression drive, not only do r2 alleles remain in the population and impair homing, but they also decrease the drive frequency (female drive/r2 heterozygotes are sterile) (Figure S1).

As the cutting rate decreases with constant drive conversion, we see that the standard suppression drive is not heavily impacted (its genetic load depends heavily on the conversion rate, with germline resistance alleles having little effect), whereas the distant-site suppression drives do not lose their effectiveness until the total cut rate falls to 0.6 (Figure 4 and Figure S2). Because the distant site drives use the total cut rate for disruption of the female fertility gene target, this total cut rate largely determines the suppressive power rather than drive conversion. Though the distant-site haplosufficient rescue drive is slower to spread due to reduced ability to remove r2 alleles, they both eventually reach the same equilibrium population suppression (Figure S2).

For each gene drive, we determined the complete suppression success rate and the genetic load (Figure 5). The genetic load is the suppressive power of a drive, defined as the reduction in reproduction of the population compared to a wild-type population of the same size(14). In Figure 5A, we see that the standard suppression drive has the smallest area of population elimination success, followed by the haplosufficient two-target drive, and then the haplolethal two-target drive. The standard suppression drive requires a high drive conversion rate to eliminate the population, whereas both two-target drives rely mostly on the cut rate alone. The same pattern is visible in the genetic load in Figure 5B, where the genetic load of the standard suppression drive relies on the conversion rate only, whereas both two-target drives rely almost entirely on the cut rate for their genetic load.

**Figure 5.**
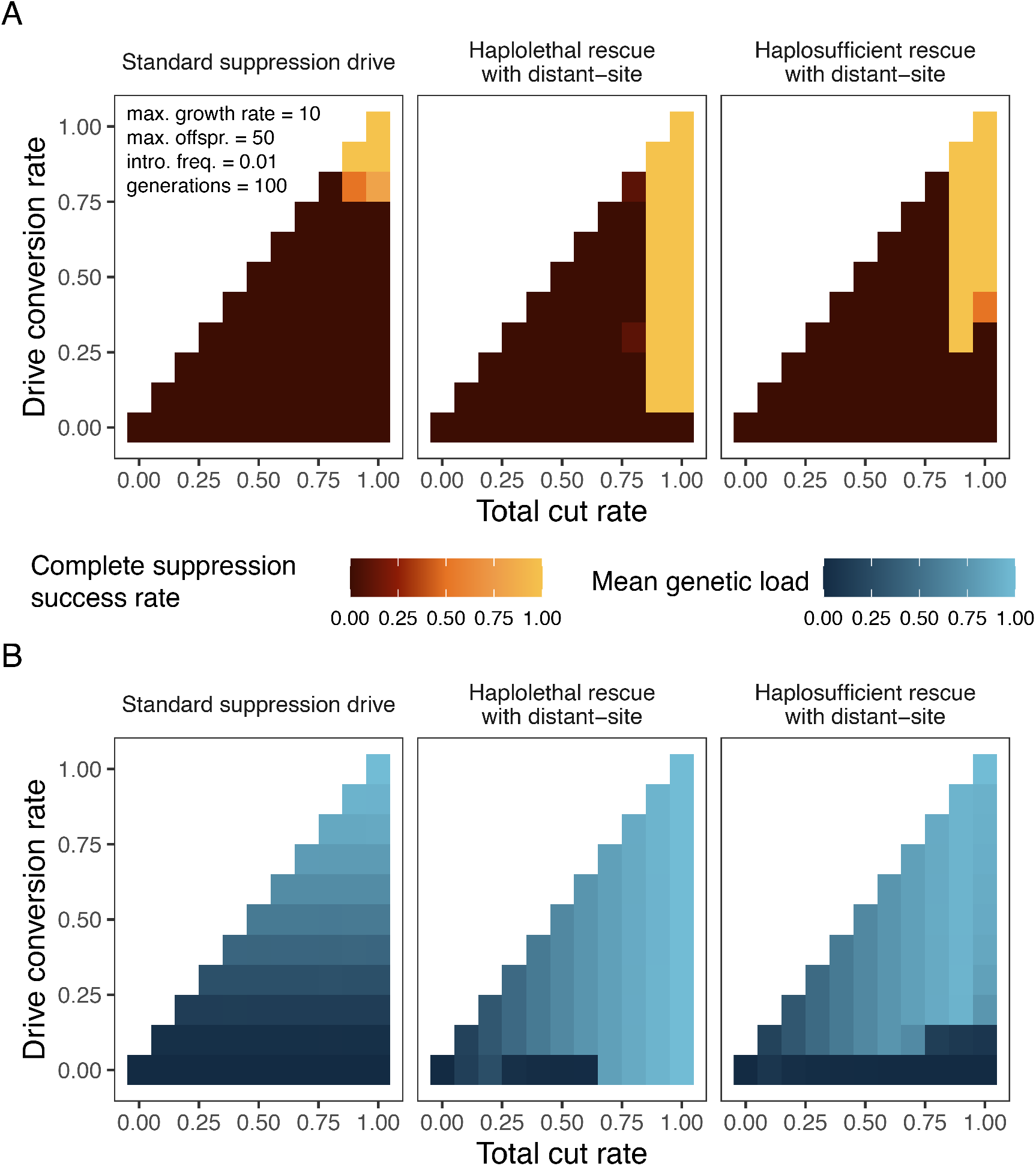
Drive performance under varying cut and conversion rates. **A)** Complete suppression success rate and **B)** mean genetic load with variable total cut and drive conversion rates. The complete suppression success rate is calculated as the number of repetitions in which population elimination occurs within 100 generations after drive introduction divided by the total number of repetitions. For each combination of parameters, we ran 10 model repetitions.

The haplosufficient rescue drive shows some more complex dynamics in some areas of parameter space in Figure 5B. When the cut rate is 1 but the drive conversion is low, the genetic load is reduced because the drive itself is not able reach high frequency due to r2 alleles are blocking its progress (Figure S3). The drive spreads best with the highest genetic load when the total cut rate is somewhat below 1. At the same time, high cut rates are still necessary at the distant site to achieve population complete suppression (Figure S3). Another notable dynamic for both two-target drives in Figure 5B is the bottom row, where drive conversion is 0. Here, the two two-target drives become identical to a version of two toxin-antidote drives previously described called a TADE (Toxin Antidote Dominant Embryo) suppression drive and a TARE (Toxin Antidote Recessive Embryo) suppression drive(41). These drives show density-dependent dynamics, so their ability to increase in frequency depends on their frequency as well as their total cut rate, hence the lack of suppression success for the haplolethal rescue drive due to our small release size. In the case of the haplolethal two-target drive, due to the additional disruption of the distant-site female fertility gene, a cut rate of at least 0.7 is necessary for the drive to increase in frequency. The TARE-like suppression drive is not able to reach a high genetic load in the first place(41). Interestingly, at very low cut rates, we observe that the two-target drives are able to remain in the population long enough for the distant-site to be disrupted up to a certain frequency, after which the gene drive can sometimes induce a small genetic load during the simulation.

#### A.2. Embryo cut rate and somatic expression effects on fitness

Two other important determinants of drive success are the embryo cut rate and fitness costs in heterozygous females based on the disruption of wild-type alleles when they are needed for fertility(25, 29). Embryo cutting occurs when Cas9 and gRNAs are maternally deposited into the embryo post-fertilisation. Any wild-type alleles (at the drive or distant target site), especially paternal wild-type alleles, can be cleaved, which always results in resistance allele formation at this stage (since the window for homology-directed repair is over). This process impairs the drive’s spread much more than germline resistance alleles because female drive progeny will be sterile, and male drive progeny will be unable to perform drive conversion. Similarly, undesired Cas9 expression in somatic cells, regardless of whether it results in drive conversion or resistance allele formation, will disrupt the wild-type alleles needed for female fertility, at least in some cells. Depending on where and when the fertility target gene is needed, even necessary germline expression could induce a similar fitness cost in female drive heterozygotes (or any female with at least one drive allele and at least one wildtype fertility target gene).

To assess the effects of embryo resistance and fitness cost under moderate drive conversion, we model a total cut rate of 0.9 and drive conversion rate of 0.5. Varying these two parameters over their full range (Figure 6A), we see that the haplolethal two-target drive is sensitive to high embryo cut rates, whereas the haplosufficient two-target drive is not strongly affected by this. Both are affected negatively by female heterozygote fitness costs, which prevent success more easily for the weaker haplosufficient drive. Looking at genetic load (Figure 6B), the standard suppression drive starts out weak, with increases in the embryo cut rate having roughly half the effect of reduction of female fitness. The two-target drives have a larger area of high genetic load compared to the area of complete suppression success, representing areas where the drive is still able to increase and eventually reach higher genetic load, but not within the time frame of the normal simulations. Higher initial release frequencies could still allow these drives to succeed in a shorter time frame.

**Figure 6.**
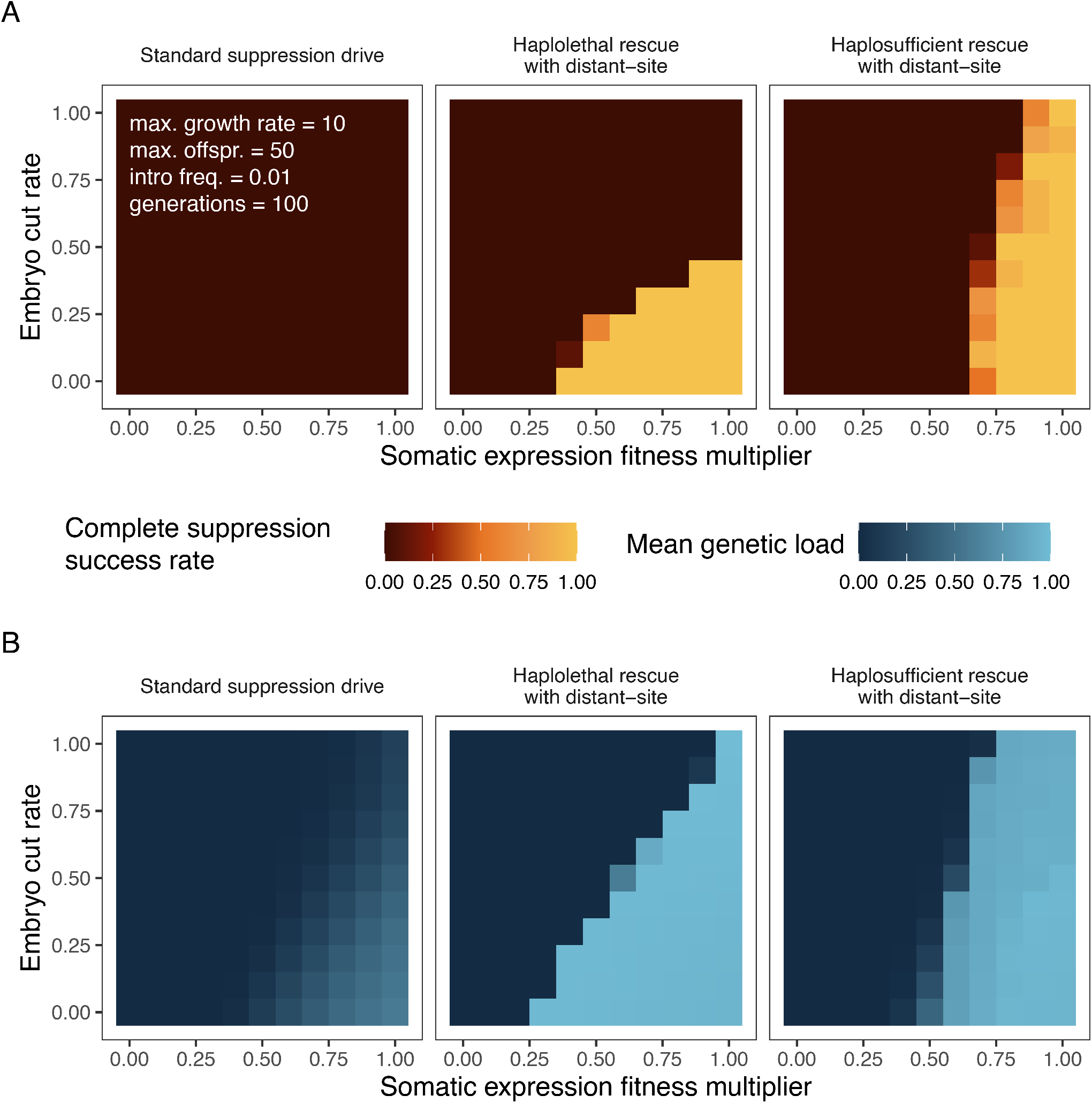
Drive performance under varying somatic and embryonic drive activity. **A)** Complete suppression success rate and **B)** mean genetic load under variable female fitness in drive heterozygotes from somatic expression fertility effects and variable embryo cut rates in the progeny of drive females. The complete suppression success rate is calculated as the fraction of simulations in which population elimination occurs within 100 generations after drive introduction. For **A**, the introduction frequency of gene drive heterozygotes is 0.1, and the total population capacity is 100,000. The total cut rate is fixed to 0.9, and the drive conversion rate is 0.5. For each combination of parameters, we ran 10 model repetitions.

To further show drive dynamics besides the eventual genetic load, Figure S5 shows population size over time after the gene drive introduction. The standard suppression drive cannot perform well with these parameters, so it always reaches an equilibrium population size, which is higher with increasing embryo cutting and fitness cost. With no somatic fertility cost and no embryo cutting, both two-target drives suppress the population rapidly. With an increasing embryo cut rate, all drives are slower for all three drives, but the haplolethal two-target drive is especially sensitive to this, losing all suppressive power (and the ability to increase in frequency at all) when the embryo cut rate is high. This is because high embryo cutting at the drive’s site results in immediate removal of drive alleles in the progeny of females (all progeny with nonfunctional resistance alleles at this site are nonviable).

With decreasing drive female fitness from somatic Cas9 cleavage, all drives are slower to suppress the population. Eventually, the drives lose the ability to increase in frequency in the first place, with the exact point being dependent on the drive (Figure S6). With low embryo cutting, the haplolethal drive remains successful for lower female fitness values, and this trend is reversed for higher embryo cutting.

#### A.3. Functional resistance alleles

Functional r1 resistance alleles have been shown to be a major challenge for many gene drives, so we investigate how well these can be handled by the two-target drive design. We only model r1 alleles at the homing site of each drive, and not at the distant site. The rescue modification drives can suffer from r1 alleles that form due to incomplete homing, where only the rescue sequence is copied, but not the rest of the gene drive(19). Thus, even with multiplexed gRNAs, there may be a lower limit for functional resistance for these sorts of drives. For a standard suppression drive, this is not an issue. However, while multiplexing can likely reduce functional resistance to low levels(25), this could reduce the drive conversion rate with more than 2-4 gRNAs, or with gRNAs that are further apart, both of which could cause drive failure due to lack of genetic load, thus leading to practical limits for multiplexing that may still permit functional resistance. At the distant site, multiplexing is more flexible because either homology-directed repair or end-joining can achieve the desired result. This could allow for more gRNAs that target the most highly conserved sites throughout the target gene, largely eliminating the chance to form functional resistance alleles, much like CRISPR toxin-antidote drives(42).

We compare the drives in a range of parameter space where all are always successful in the absence of r1 alleles (Figure S8). We see that at relatively low r1 rates, the standard suppression drive is the first to succumb to functional resistance allele formation in some simulations (Figure 7). This is because r1 alleles provide immediate benefit, directly allowing females to be fertile, which results in rapid increase in frequency of the r1 allele at low population density (Figure S9). Thus, if a resistance allele forms, it only needs to avoid stochastic loss for a small number of generations before it will prevent population elimination. Both two-target drives remain successful in all repetitions up to an r1 rate of 0.1. Then, the haplosufficient two-target drive fails in all repetitions, whereas the haplolethal two-target drive remains successful in most, but not all, replicates (Figure S8). In these drives, r1 alleles have only a modest advantage over the drive allele, and this is indirect. Individuals will not experience the modest female heterozygote fitness cost if they have only r1 and wild-type alleles at the drive site, and the r1 allele will not itself disrupt the female fertility gene for progeny, potentially increasing its chance of being passed on to a fertile female. However, the drive will reach high frequency (Figure S9), meaning that most r1 alleles will be together with drive alleles, limiting their advantage and thus doing little to prevent suppression, which only requires one drive allele to disrupt the distant-site female fertility gene.

**Figure 7.**
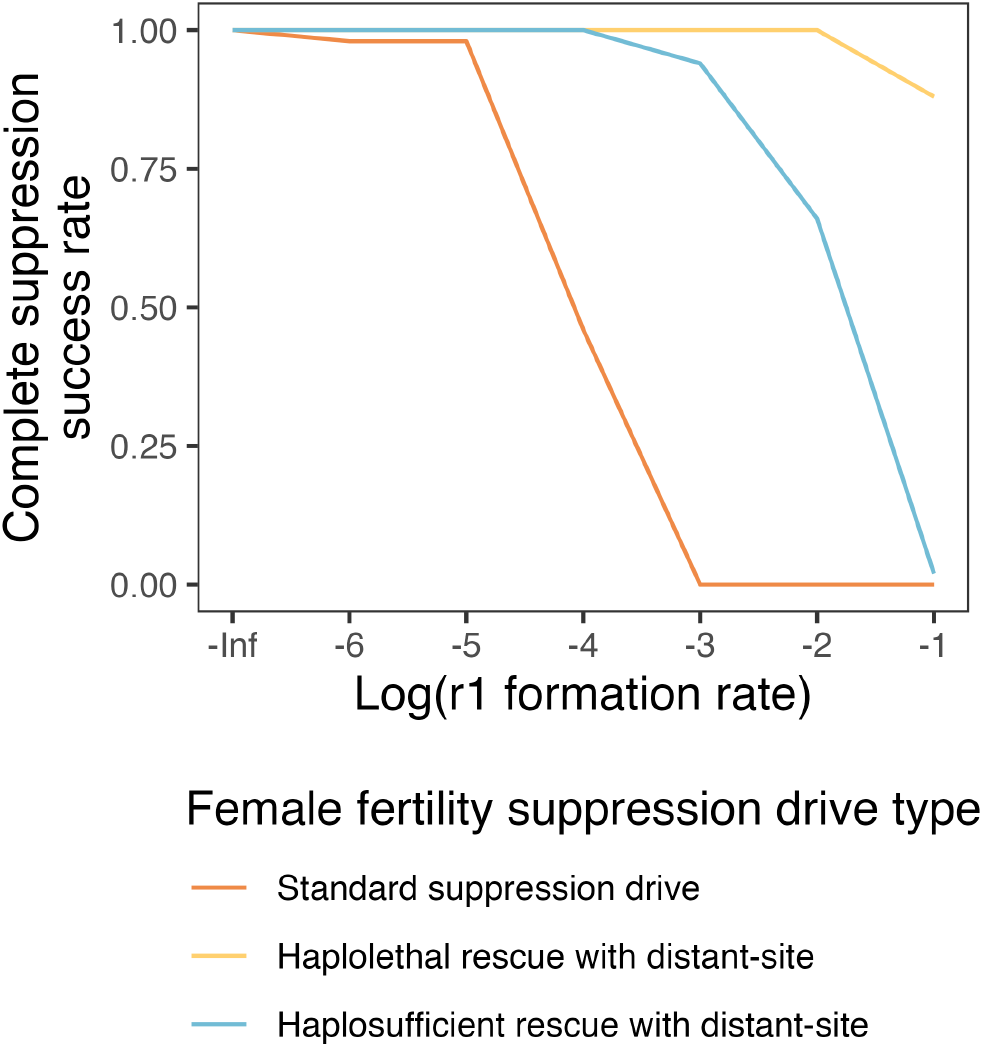
Impact of functional resistance alleles on drive performance. Successful population elimination under various functional resistance allele formation rates. The r1 formation rate is the relative rate of resistance alleles that are functional, rather than nonfunctional r2 alleles. The model is run for 100 generations after an introduction of drive heterozygotes at 0.01 frequency into a population of 100,000. To compare the drives where they were all successful in the absence of functional resistance alleles, we used a total cut rate 1.0, drive conversion rate 0.9, somatic expression female fertility fitness effect of 0.9, and an embryo cut rate of 0.1 in the progeny of female drive carriers. For each combination of parameters, we ran 50 model repetitions.

#### A.4. gRNA saturation

Because the two-target drives need double the amount of gRNAs, the drive as a whole may suffer from additional gRNA saturation compared to a standard homing suppression drive. This means that the cutting efficiency of each individual gRNA is reduced due to limited amount of Cas9 protein. Gene drives using multiplexing to avoid r1 alleles already face this challenge(19), which would be amplified in two-target drives. We do not model multiplexing explicitly, but instead assume that an equal number of gRNAs will be used for the homing site and the distant site, thus allowing us to reduce the cutting efficiency at each site proportionally. Here, we calculated the genetic load for each drive with various cut rates and gRNA saturation factors (Figure 8). The gRNA saturation factor is the relative Cas9 activity level with unlimited gRNAs (spread equally between all the gRNAs), with “1” being the activity in the presence of a single gRNA(19). A saturation factor of 1 thus means that the total cut rate is immediately split between any number of gRNAs, while a rate of infinity (our default in previous simulations) means that the cut rate at each gRNA target is the same as the cut rate of a 1-gRNA system.

**Figure 8.**
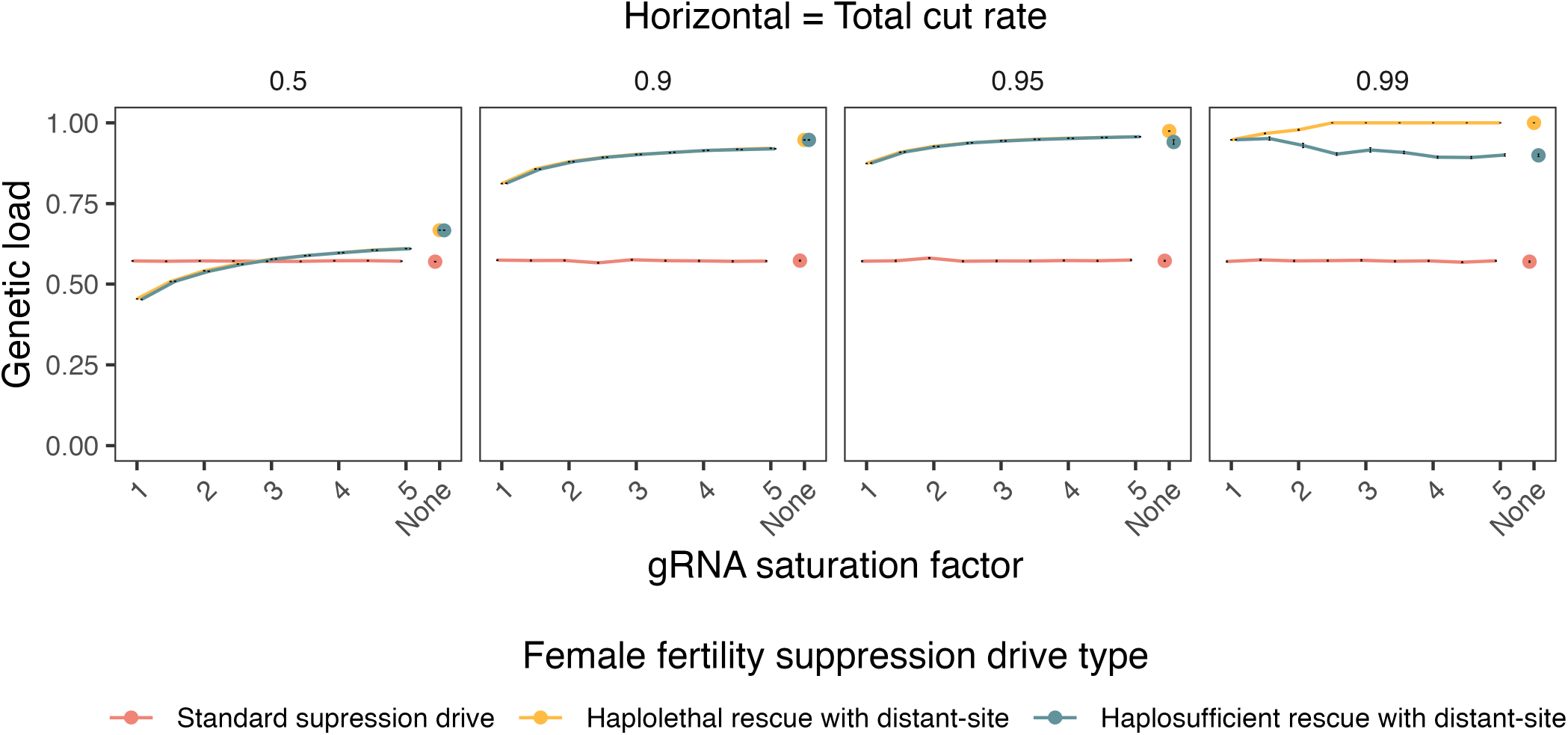
Mean genetic load with variable gRNA saturation factor and total cut rates. The gRNA saturation factor is modelled as relative Cas9 activity with unlimited gRNAs compared to the activity with a single gRNA. We assume that the distant site target gene drives have double the amount of gRNAs compared to a standard suppression drive and that the cut rates are equally reduced at both sites (the two sites are assumed to have the same number of gRNAs). The specified total cut rate is the rate for unlimited gRNA saturation factor. We used a drive conversion rate 0.5, somatic expression female fertility effect of 1, and embryo cut rate of 0. For each combination of parameters, we ran 10 model repetitions.

The standard suppression drive is not impacted by this model of gRNA saturation because this drive only targets a single site (it can be thought of as the “baseline” in this scenario, even if it would have multiple gRNAs in practice) (Figure 8). The two-target drives are both impacted by gRNA saturation, which reduces the genetic load a moderate amount if the gRNA saturation factor is low. This reduction is low if the total cut rate is high (Figure S12). For the haplosufficient drive with a cut rate of 0.99, gRNA saturation actually has a positive effect on the drive because its optimal cut rate is somewhat lower than 0.99 (Figure S13). The current best estimate of the gRNA saturation factor is 1.5(19), though this could potentially vary substantially between systems. Overall, gRNA saturation is potentially problematic, though the two-target drives would still usually be expected to have higher power than a standard suppression drive, especially when the total cut rate is high.

### B. Proof of concept of two-part gene drive in *Drosophila melanogaster*

We constructed the two-target gene drive design by reusing two previously built gene drives, a standard female fertility suppression drive and a haplolethal modification drive, and combining them(25, 36). We show that the gene drive has moderate drive conversion in both male and female drive heterozygotes, and that cutting at the distant site is highly efficient. However, we also show that drive heterozygous females suffer a significant fertility cost that usually prevented the gene drive from eliminating cage populations.

#### B.1. Drive crosses

To demonstrate a proof-of-principle for this novel drive design, we developed a suppression drive in *D. melanogaster*, with the drive construct integrated in the haplolethal gene *RpL35A*. The distant target site is the haplosufficient female fertility gene *yellow-g* (Figure 9A). This drive was constructed based on another successful drive reported previously(36), by adding extra gRNAs targeting *yellow-g* from a previous suppression drive(25) to convert the original modification drive into a suppression drive.

**Figure 9.**
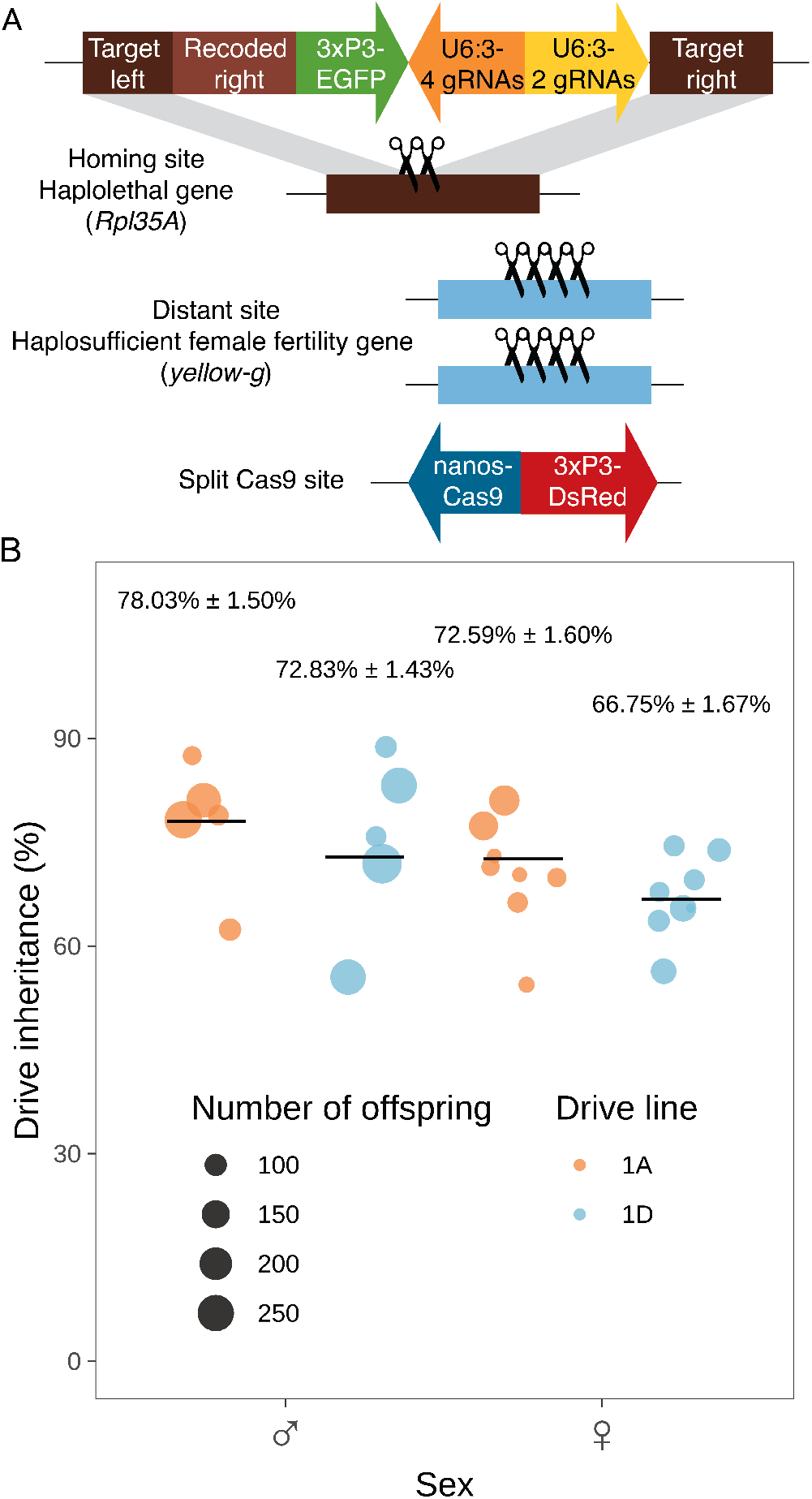
Drive construct and inheritance rates. **A)** The drive allele is inserted into the haplolethal gene *RpL35A* and contains a recoded version as rescue to avoid gRNA cleavage. It is expected that only offspring inheriting two functional copies of *RpL35A* (which can be either a drive or wild-type allele) can survive. The two blue lightning bolts show the gRNA sites for cleaving *RpL35A* and homing, while the four brown lightning bolts show the gRNA sites for disrupting *yellow-g*, a haplosufficient female fertility gene located at a distant site. **B)** The drive inheritance rate indicates the percentage of offspring with EGFP fluorescence from the cross between heterozygotes (containing one copy of drive and one copy of Cas9) and w1118 flies (without drive and Cas9). The size of each dot represents the total number of offspring. The inheritance rate of paternal drive is significantly higher than that of maternal drive in both line 1A (Fisher’s exact test, *p* = 0.0153) and line 1D (*p* = 0.0057).

Two independent EGFP-marked drive lines (named as line 1A and line 1D) were generated. These drive lines were respectively crossed to a DsRed-marked *nanos*-Cas9 line to produce heterozygous individuals carrying one copy of the drive and one copy of the Cas9 allele, after which these heterozygotes were crossed to w1118 flies for assessing drive efficiency. The drive inheritance rates of Line 1A were 78% for drive males and 73% for drive females, while the inheritance rates of Line 1D were 73% for drive males and 67% for drive females, significantly higher than the 50% expected with Mendelian inheritance (Binomial exact test, *p* < 0.0001**** for all four comparisons) (Figure 9B, Supplementary Data Set S1). The drive inheritance rates were much lower than the original haplolethal homing drive, where drive inheritance rates for both male and female heterozygotes were 91%. This difference is likely caused by gRNA saturation, meaning that the two gRNAs of the homing drive had lower cut rates because they shared the same amount of Cas9 with the additional four gRNAs targeting *yellow-g*. Besides the difference between males and females, the two gRNA lines performed slightly differently in drive conversion rate as well. To find the cause of this difference, Sanger sequencing was conducted to compare their genomic structures. Sequencing results showed that line 1D contained the full construct as expected, while line 1A showed recombination inside the construct (flipping of the DNA between the U6 promoters for the gRNA cassettes targeting *yellow-g* and *RpL35A*).

#### B.2. Cage experiments

To characterize the population suppression activity of this drive, cage experiments were set up by mixing drive carriers (heterozygous for the drive allele and homozygous for Cas9) and a small fraction of Cas9 homozygous flies. These cages were followed for several discrete generations, measuring drive carrier frequency by phenotyping all adults. Though there was some stochasticity in population size due to the single bottle nature of the cages, the dynamic of the drive was clear. Drive carrier proportions of most cages quickly decreased within five generations (Figure 10A). The population size for the two lower frequency releases was unaffected, and for one of the high release frequency cages, the population size recovered after an initial reduction (Figure 10B, Supplementary Data Set S2). However, the other high frequency release cage showed complete population suppression in the second generation even though drive carrier frequency was declining. In the suppressed population, the five remaining flies all happened to be females, whereas this was not the case in the population that recovered. There, the five flies in that generation consisted of four females (one of which lacked the drive) and one male (also without the drive), allowing the population to grow immensely in the next generation. This result implies that the distant-site suppression drive was functional, though overall efficiency was low due to drive fitness costs outweighing biased drive inheritance. It also confirms that near-fixation of a gene drive will not be enough to suppress a population of *D. melanogaster*, given their ability to produce many offspring from just a few remaining individuals under the conditions of these cages.

**Figure 10.**
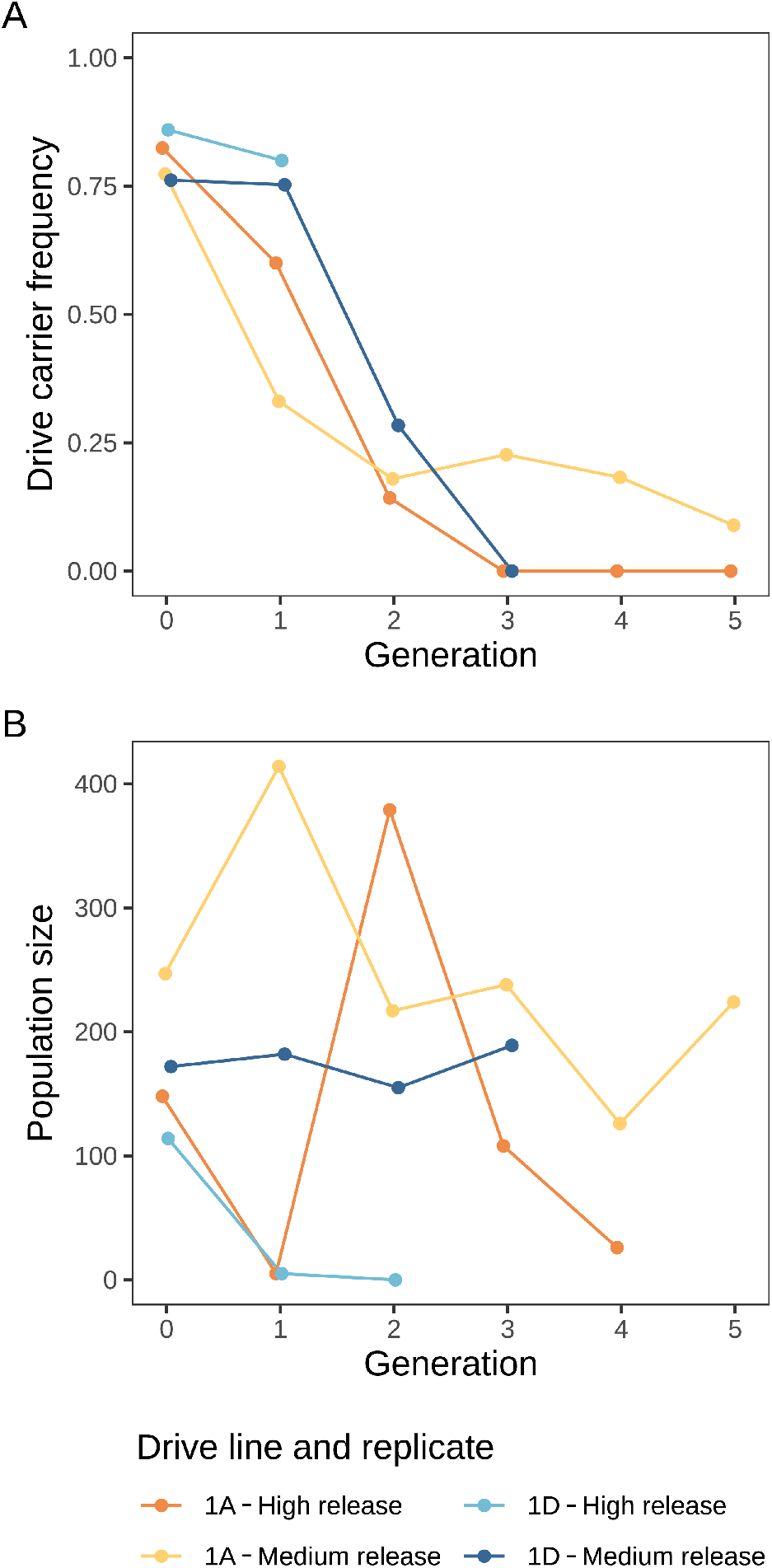
Small cage experiments. Heterozygous flies carrying one copy of the drive allele and two copies of Cas9 were used to generate progeny for “generation zero” of four small cages (the two medium release cages also included a non-drive female that was also homozygous for Cas9). **A)** Drive carrier frequency and **B)** population size in each generation.

#### B.3. Resistance alleles and fertility

While functional resistance is a possible explanation for the rapid decline of a suppression drive from high frequency, we considered this unlikely due to the use of multiplexed gRNAs and previous studies in larger cages with similar drives finding no evidence of functional resistance(25, 36). To explore the possible reason for poor cage performance, a fertility test was conducted to assess the fitness of our drive carriers (which was found to be reduced in both drives this two-target drive is based on, but not to the extent of eliminating them from cages(25, 36)). This fitness cost is based on egg viability from drive females, which was much lower compared with drive males and controls without the drive. In non-drive females, egg viability was and perhaps also slightly lower (Figure 11, Supplementary data set S3), though this latter effect could potentially be ascribed to a batch effect due to the purported haplosufficiency of *yellow-g*. We collected 16 and 18 drive daughters from drive females (which were offspring of the above cross) of line 1A and line 1D, respectively, and crossed them to *w1118* males. 87.50% from line 1A and 77.78% from line 1D were sterile (Supplementary data set S4), which was likely caused by resistance allele formation in the embryo at the *yellow-g* target site due to maternal deposition of Cas9 and gRNAs. Indeed, these sterile females were collected for sequencing, which indicated 100% cutting in the *yellow-g* target site. Another 16 non-drive progeny from drive female and Cas9 male cross were randomly collected for sequencing, showing 87.5% (14 out of 16) cutting in *yellow-g*. These results indicate that the poor cage performance in this study is likely due to fitness cost of female drive heterozygotes together with high levels of embryo resistance allele formation at the female fertility site and some embryo resistance allele formation a the drive site, though it is currently unclear to what extent each of these factors contributes.

**Figure 11.**
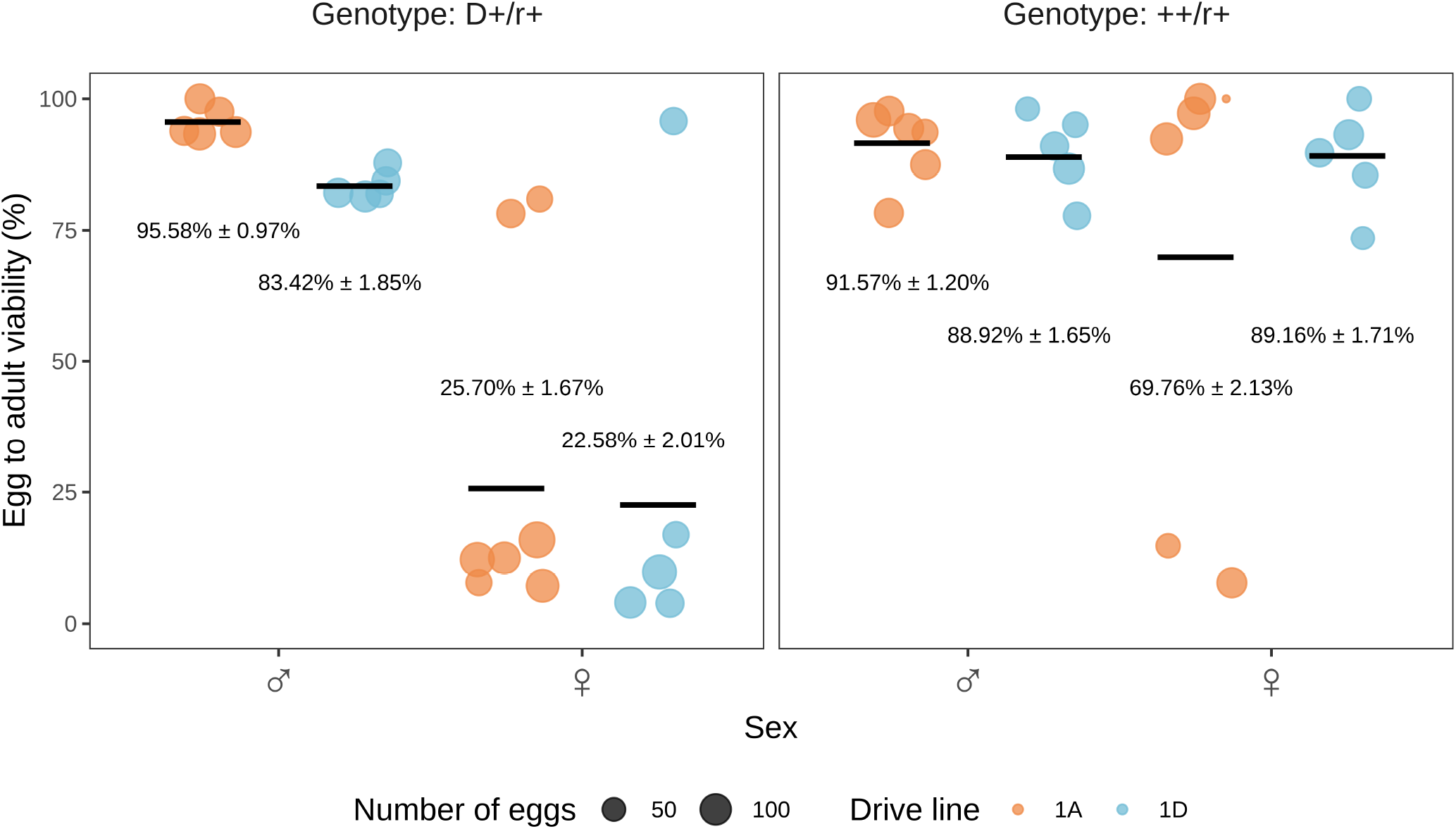
Egg viability test. The individuals tested in this experiment were all homozygous for Cas9, but different in the drive site and distant site, for which genotypes are listed at the top. “+”, “D” and “r” respectively represent wild-type allele, drive allele and nonfunctional (*yellow-g* target site) resistant allele. Note that while cut rates at the *yellow-g* site were high, it is possible that a small fraction of flies marked as having an “r” at the target site were in fact homozygous for wild-type at this site.

In addition to the egg to adult viability, we measured the drive inheritance of these offspring. The drive inheritance rate of line 1A was 67.67% ± 2.25% in drive males and 73.71% ± 3.33% in drive females, while it was 75.96% ± 2.83% in drive males and 77.75% ± 4.21% in drive females in line 1D (Figure S15). The drive inheritance rates of these flies were somewhat higher than our previous test where drive heterozygotes had only a single copy of Cas9, except for drive males of line 1A. The reason for this anomaly is unclear, particularly considering that lack of Cas9 appears to be a limiting factor in the performance of the drive compared to the original modification drive it is based off. Additionally, although drive inheritance rates in the previous test were higher in drive males than drive females, this difference was not seen in this later drive conversion test.

## Discussion

In this study, we analyzed the possibility of converting a modification to a suppression drive by addition of gRNAs targeting a female fertility or other essential gene without rescue. We found via modelling that this can substantially increase the genetic load (suppressive power) of the drive compared to a similar standard suppression drive, unless the drive conversion rate is already very high. Instead of the genetic load being mainly determined by the drive conversion rate, it is determined by the total germline cut rate in these two-target drives. We then demonstrated such a drive in the model organism *D. melanogaster*, which successfully biased its inheritance while cutting the distant female fertility gene target at high rates. However, fitness costs in drive heterozygous females were quite high for this drive, which prevented success in most cage replicates despite high release frequency. Nevertheless, two-target drive systems may be quite desirable tools for suppression of pests where high drive conversion is difficult to achieve but not high cut rates. Such high drive conversion has only been consistently achieved in *Anopheles* mosquitoes, while several other species have had substantially lower or inconsistent drive conversion, such as *Drosophila, Aedes*, and mice(20, 24, 25, 30, 32, 37). In these, relatively high cut rates have often been achieved when drive conversion is low, which would be necessary for the two-target drive design to work in these species.

In the two-target suppression drive, population suppression is largely decoupled from drive frequency increase. This is in contrast to the standard homing suppression rive, where the drive allele itself is mainly responsible for disrupting the suppression target. Thus, the two-target drive can keep increasing in frequency and eventually cut most female fertility target alleles. The standard suppression drive reaches an equilibrium that allows many wild-type alleles to remain, mostly accounting for its genetic load. Though the two-target suppression drive allows for higher suppressive power, it still needs to be able to increase in frequency in the first place. Thus, drive conversion must be sufficiently high to overcome fitness costs and embryo resistance (the latter of which is mostly important at the distant site, though it could have a substantial negative effect at the drive site if the drive targets a haplolethal gene there). These performance parameters are still highly dependent on the Cas9 promoter and to a lesser extent, the target gene. For the target gene, additional possibilities are available for two-target drives because they can keep high genetic load even when targeting X-linked female fertility genes.

While our modeling results are promising for two-target drives that lack high drive conversion rates, it should be noted that drive conversion is still essential for determining the speed at which the drive’s frequency will increase, which determines the time to population elimination. Though this system can increase the genetic load, such drives will still take longer to eliminate a population than a drive with high drive conversion efficiency (regardless of whether such a drive is a standard suppression drive or a two-target system). In more complex environments such as two-dimensional continuous space, this slowed rate of increase could potentially result in failed suppression due to the “chasing” effect, where wild type individuals migrate back into an area previously cleared by the gene drive, resulting in cycles of suppression and reestablishment(12). However, this same study included a powerful but slow drive that avoided this effect, which may be analogous to a two-target system. Additional modeling is thus needed to explore the outcomes of two-target drives in complex scenarios.

In our experimental demonstration, the constructs functioned generally as expected, performing drive conversion while cutting the distant site at high rates. However, drive efficiency at the homing site was considerably lower than the original modification rescue drive(36). Because we used the same Cas9 and gRNAs, the reason for this was likely Cas9 saturation from the additional gRNAs targeting the distant site. With only two gRNAs for the homing site and four gRNAs for the distant site (both selected based on previously available constructs(25, 36)), the cut rate at the homing site would be expected to decrease more than at the distant site (in our model we assume that the same number of gRNAs target each site), especially because the cut rate for the original modification drive was not high in the first place(36), at least compared to other published drives in *D. melanogaster*. These issues could be more readily avoided using entirely new designs with balanced numbers of gRNAs.

Arguably more problematic, the high fitness cost in female heterozygotes prevented success in population cages, and embryo resistance at both the drive site and the distant-target site further contributed to drive failure. This fitness cost was expected to an extent because our two-target drive is based on a suppression drive showing reduced fitness as well(25). In the haplolethal modification drive, drive carrier males showed no significant reduction in offspring egg-to-pupa survival, but females did show a significant effect(36). This fitness reduction would represent an addition fitness cost in our two target drive and is likely from nonfunctional resistance alleles rendering offspring nonviable. If formed in the germline, these alleles would not have significant negative effects, but if formed in the embryo, it could further contribute to reduced drive success. In the earlier standard suppression drive targeting *yellow-g*, at first, offspring from neither male or female drive parents showed a significant reduction in the egg-to-pupa survival rate, but later crosses with drier food conditions did show a significant reduction in egg-to-pupa survival rate, and inference from cage experiments showed even higher fitness costs. This fitness cost likely indicates either leaky somatic expression or disruption of *yellow-g* too early in the germline where it still might need to function. Together, these factors could all contribute to the more severe fitness cost for female drive carriers with our two-target drive, though we currently do not know what factor contributes to what extent. In the original suppression drive, offspring of drive carrier females were found to be sterile at high rates. This infertility indicates significant embryo activity, which also had a similar negative effect on the two-target drive.

In the two-target drive, gRNA saturation could have slightly ameliorated these effects due to the presence of more gRNAs and cut rates being slightly reduced, but the different genomic site may change this, and reduction in drive conversion efficiency for similar reasons would likely outweigh any benefits.

All of these fitness costs will likely be similar for two-target drives targeting different genes using *nanos*-Cas9, with the exception of *yellow-g* potentially being required in the early germline. Therefore, finding promoters that are exclusively expressed in the (late) germline is essential for future development of two-target drives (and any suppression drive for that matter), and promoters with little to no embryo activity are also preferred. In the standard suppression drive, however, the higher drive conversion rate (and lack of embryo cutting at the haplolethal target) still allowed it to reach a moderate equilibrium instead of declining to zero. If the two-target drive in our study had similar or even somewhat lower drive conversion, our model indicates that it would have been able to increase in frequency in the population and eventually eliminate it with high genetic load. Even in current form, our drive could potentially be successful with repeated releases, analogous to a more powerful form of sterile insect technique, which is far less powerful than a gene drive but still potentially useful in several situations due to its self-limiting nature(43).

Overall, we have shown that for scenarios of low to moderate drive conversion and high total germline cut rates, two-target drives may offer substantially increased suppressive power. With further efforts to find better Cas9 promoters and target gene combinations with lower fitness costs, this unique drive design may unlock the potential for strong population suppression in many scenarios.

## ACKNOWLEDGEMENTS

This study was supported by the Graduate School for Production Ecology & Resource Conservation (PE&RC) in the Netherlands, laboratory startup funds from Peking University, the Center for Life Sciences, the Li Ge Zhao Ning Life Science Youth Research Fund, and grants from the National Science Foundation of China (32302455, 32270672, and Overseas Youth).

## Supplementary Materials

### A. Supplementary methods

#### A.1. Drive types

We have modelled 10 different types of homing suppression drives. The first three in the list below are shown in the main results. Modelling for the rest of the drives are shown here in the Supplementary Materials.

1. Drive with female fertility target
2. Haplolethal rescue drive with distant-site female fertility target
3. Haplosufficient rescue drive with distant-site female fertility target
4. X-linked female fertility target
5. Haplolethal rescue drive with distant-site X-linked female fertility target
6. Haplosufficient rescue drive with distant-site X-linked female fertility target
7. Haplosufficient but essential target
8. Haplolethal rescue drive with distant-site haplosufficient but essential target
9. Haplosufficient rescue drive with distant-site haplosufficient but essential target
10. Standard female fertility drive with distant-site female fertility target

The methods are all described in the main text, except for one detail for drives number 7, 8 and 9. Since these drives target an essential gene instead of a female fertility gene to achieve population suppression, we have implemented the somatic expression fitness cost as an embryo viability effect rather than a female fertility effect. The formula remains the same as in Formula 4.

### B. Supplementary results

#### B.1. Additional drive types

Besides our three main types of drives, we have modelled seven more (see Section A.1). These seven include various drives with X-linked targets and viability targets (that affect both sexes, rather than only fecundity and only in females), plus one drive that does not include a rescue element and instead operates similar to a standard suppression drive at its homing site (while still targeting a second distant female fertility gene).

First, although the standard drive loses suppressive power more quickly when the female fertility gene is X-linked(44), the two distant-site drives show no significant decrease in suppressive power when the main drive is on the autosome and the distant target is X-linked (Figure S4). This difference is that homing can still occur in both sexes in this situation, allowing the drive to increase in frequency more rapidly. Because there are many genes located on the X chromosome that are differentially expressed between males and females and might play a role in female fertility(45, 46), this possibility could increase the number of potential target genes while still retaining high drive efficiency.

Second, a standard gene drive that targets a both-sex viability genes requires higher performance to achieve the same success as a drive targeting female fertility (Figure S4).

Distant-site targeting of such a gene does increase the genetic load, but performance is still weaker than a female fertility target, especially if the rescue element is for a haplosufficient gene. In this latter variant, higher total cut rates beyond the level of the drive conversion rate will first increase but eventually reduce the drive’s genetic load, even though it still offers superior performance to a standard drive targeting a both-sex viability gene.

Third, the standard female fertility drive with an extra distant-site female fertility target has an area of complete suppression success where none of the other drives do with lower total cut rate (because cutting at both targets now contributes to genetic load), though the drive conversion rate still needs to be relatively high for success. This drive always performs worse with higher total cut rate if the drive conversion rate is fixed.

We further show that the two-target drives that target female fertility are more robust against embryo cutting and somatic expression fitness costs than their standard drive counterparts (Figure S7), although it must be noted that some drives (including the standard X-linked drive and standard viability drive) are already unable to achieve any genetic load under the chosen parameters.

Modelling functional resistance, we show that two drives appear superior to all others, still achieving complete population suppression in most replicates: the haplolethal rescue drive with distant site female fertility target, and the same drive with an X-linked fertility target (Figure S10 and S11). Though some of the modelled drives are unable to achieve complete population suppression even in the absence of functional resistance alleles, we show that the two-target drives are consistently better than their single-target counterparts drives targeting the same gene in preventing functional resistance alleles from rescuing the population.

Finally, we find that all two-target drives are similarly impacted by gRNA saturation (Figure S14), with the notable exception of the haplosufficient rescue drive with a distant-site viability target. This drive benefits most from the conversion rate being close to the cut rate (Figure S4), so gRNA saturation can benefit the drive in some cases. For the same reason, though most drives show a higher genetic load with a higher cut rate, the standard female fertility drive with distant-site female fertility target shows a somewhat reversed trend.

### C. Supplementary figures

**Figure S1.**
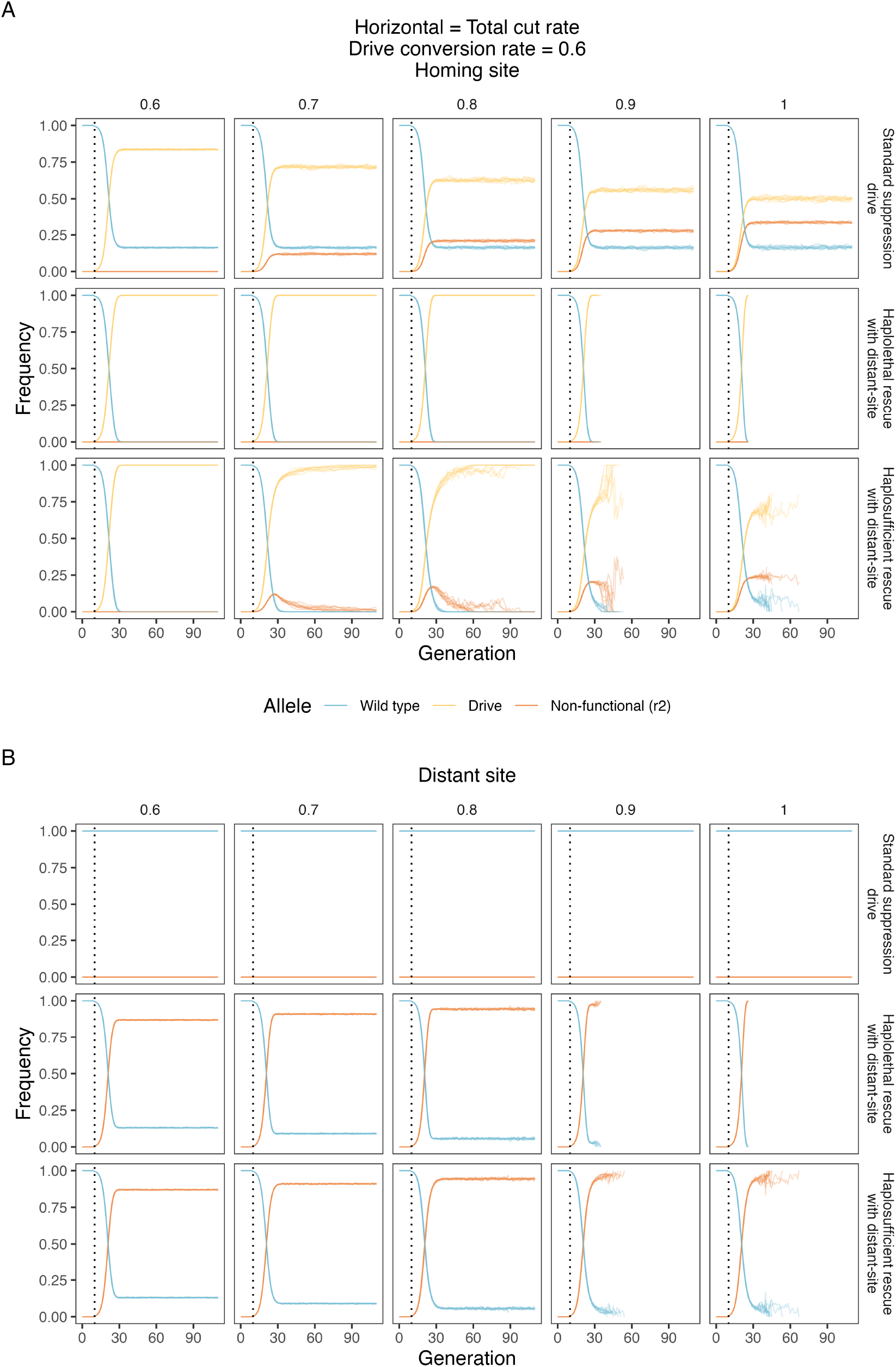
Example allele frequency trajectories. **A)**homing site and **B)** distant site allele frequencies under a conversion rate of 0.6 and varying total cut rate. The gene drive is introduced after generation 10 (the dotted line). The introduction frequency of gene drive heterozygotes is 0.01, and the total population size is 100,000. Note that both drive conversion rate and total cut rate are absolute rates, so the drive conversion rate can never be higher than the total cut rate. For each combination of parameters, we ran 10 model repetitions for each drive.

**Figure S2.**
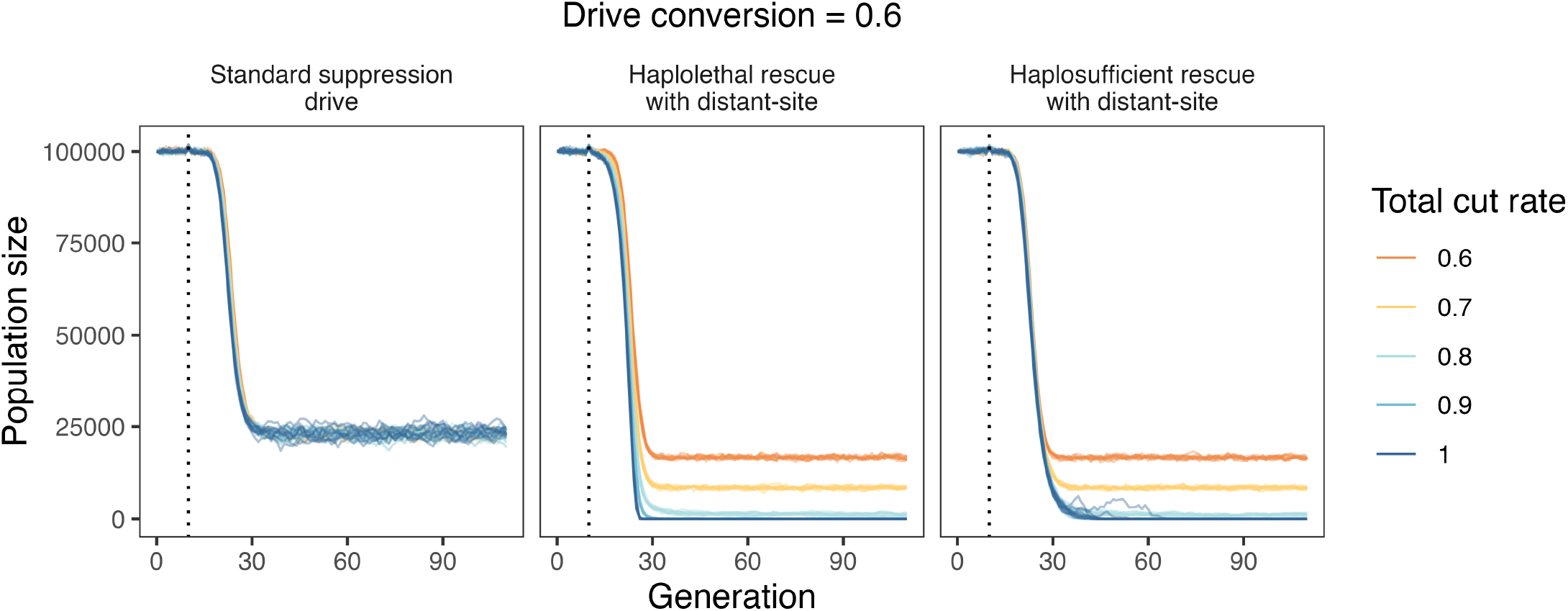
Population suppression under varying cut rates. Population size under a conversion rate of 0.6 and varying cut rates. The gene drive is introduced after generation 10 (the dotted line). The introduction frequency of gene drive heterozygotes is 0.01, and the total population size is 100,000. Note that both drive conversion rate and total cut rate are absolute rates, so the drive conversion rate can never be higher than the total cut rate. For each combination of parameters, we ran 10 model repetitions for each drive.

**Figure S3.**
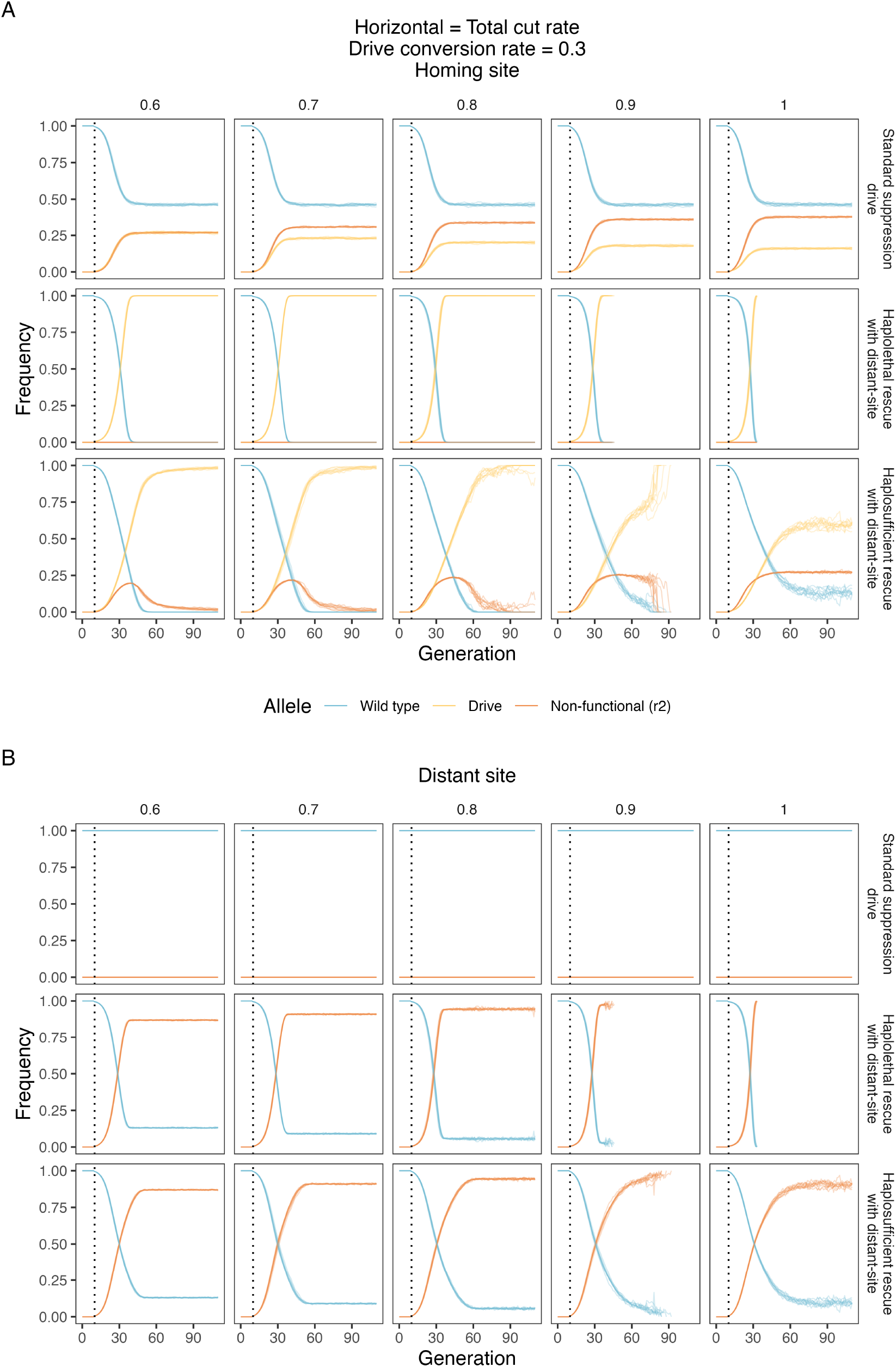
Example allele frequency trajectories. **A)**homing site and **B)** distant site allele frequencies under a conversion rate of 0.3 and varying total cut rate. The gene drive is introduced after generation 10 (the dotted line). The introduction frequency of gene drive heterozygotes is 0.01, and the total population size is 100,000. Note that both drive conversion rate and total cut rate are absolute rates, so the drive conversion rate can never be higher than the total cut rate. For each combination of parameters, we ran 10 model repetitions for each drive.

**Figure S4.**
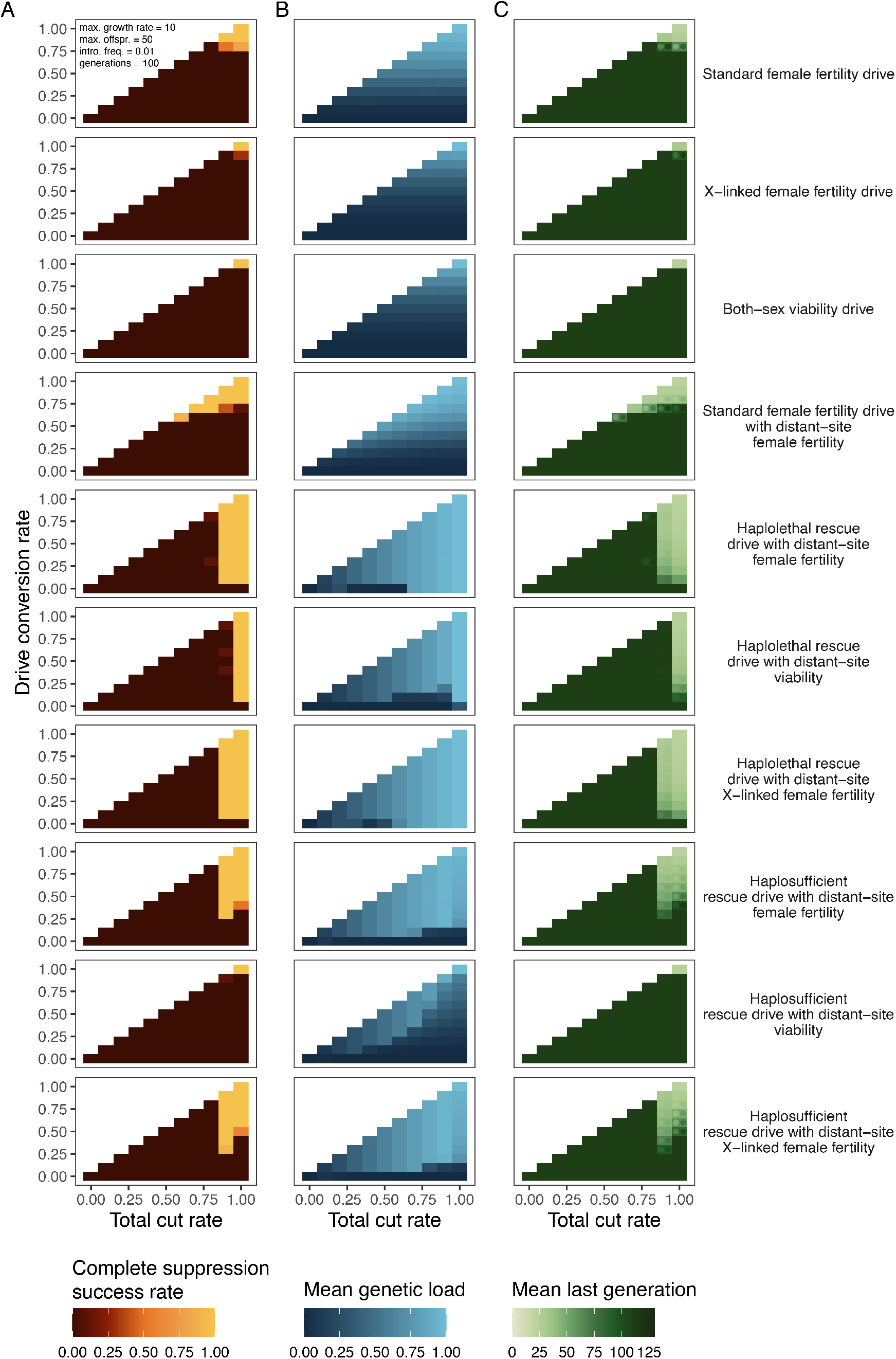
Drive performance heatmaps for additional drive variants. **A)**Population elimination success rate, **B)** mean genetic load, and **C)** number of generations before population elimination for all ten gene drive types under various cut and conversion rates. In **C**, the coloured circles in the heatmap show the 95% confidence interval around the mean. The complete suppression success rate is calculated as the number of repetitions in which complete suppression occurs divided by the total number of repetitions. Because the model stops at 111 generations, a last generation of 111 means that suppression was not successful. For each combination of parameters, we ran 10 model repetitions.

**Figure S5.**
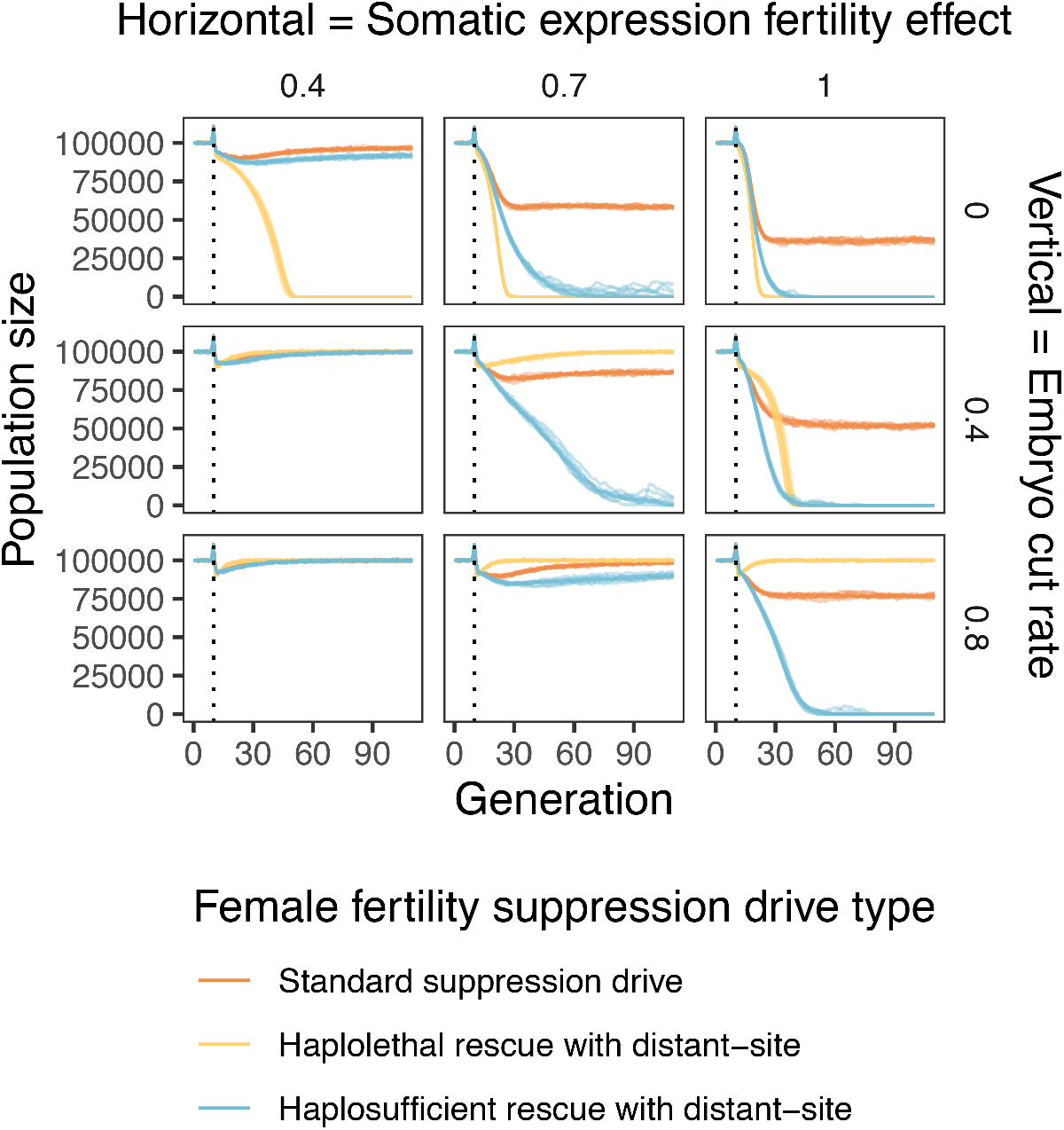
Population trajectories under variable somatic and embryonic drive activity. female fitness in drive heterozygotes from somatic expression fertility effects and variable embryo cut rates in the progeny of drive females. The gene drive is introduced after generation 10 (the dotted line), and the model is run for a further 100 generations. The introduction frequency of gene drive heterozygotes is 0.01, and the total population capacity is 100,000. The total cut rate is fixed to 0.9, and the drive conversion rate is 0.5. For the somatic expression fertility effect, 1 means there is no reduction in fertility, and 0 means females are completely sterile when the gene drive is present. For each combination of parameters, we ran 10 model repetitions.

**Figure S6.**
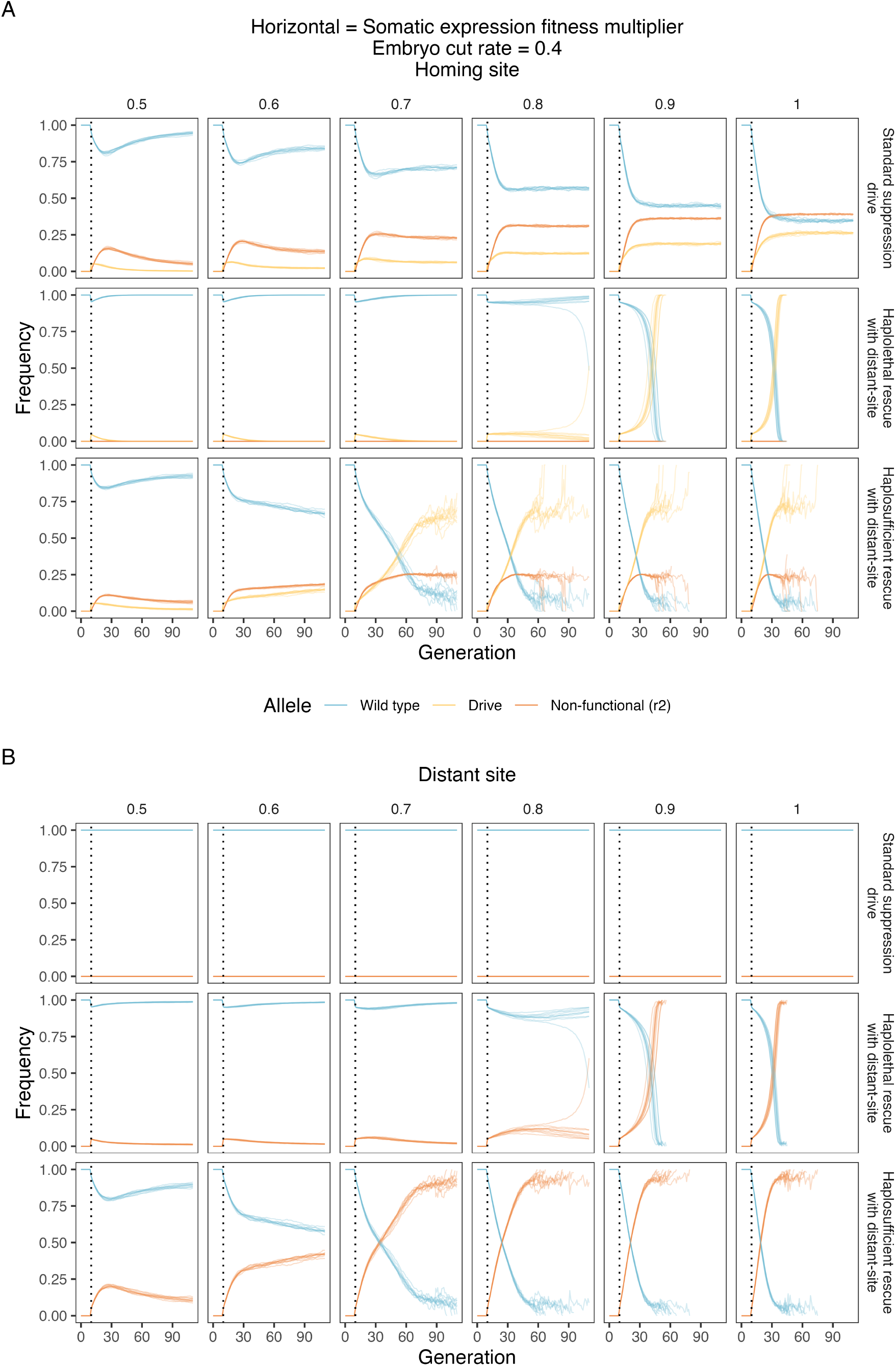
Example allele frequency trajectories. **A)**homing site and **B)** distant site allele frequencies under an embryo cut rate of 0.4 and varying somatic expression fitness multipliers. The gene drive is introduced after generation 10 (the dotted line). The introduction frequency of gene drive heterozygotes is 0.01, and the total population size is 100,000. For each combination of parameters, we ran 10 model repetitions for each drive.

**Figure S7.**
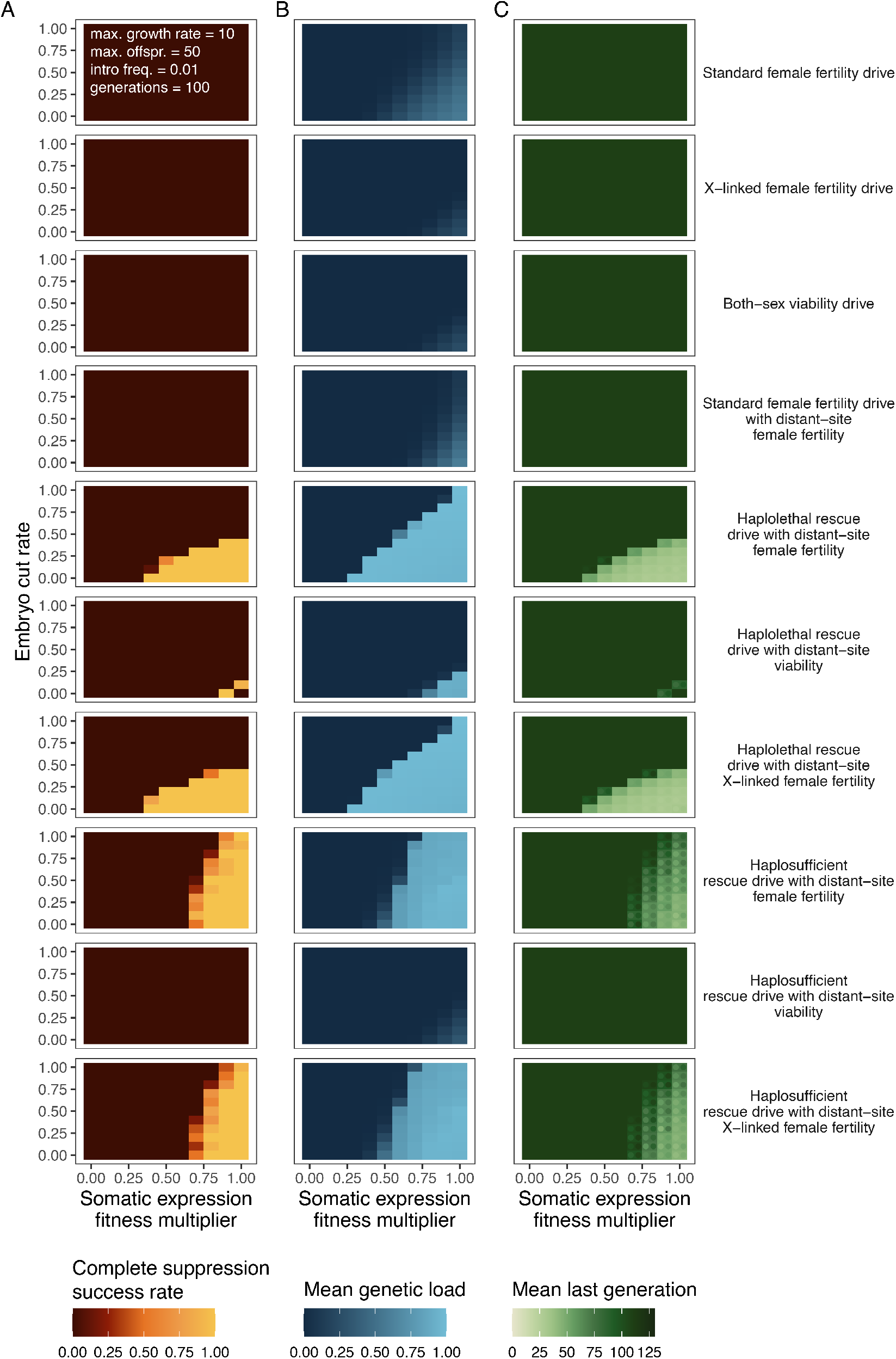
Drive performance heatmaps for additional drive variants. **A)**Population elimination success rate, **B)** mean genetic load, and **C)** number of generations before population elimination for all ten gene drive types under various embryo cut rates and somatic expression multipliers. In **C**, the coloured circles in the heatmap show the 95% confidence interval around the mean. The complete suppression success rate is calculated as the number of repetitions in which complete suppression occurs divided by the total number of repetitions. Because the model stops at 111 generations, a last generation of 111 means that suppression was not successful. For each combination of parameters, we ran 10 model repetitions.

**Figure S8.**
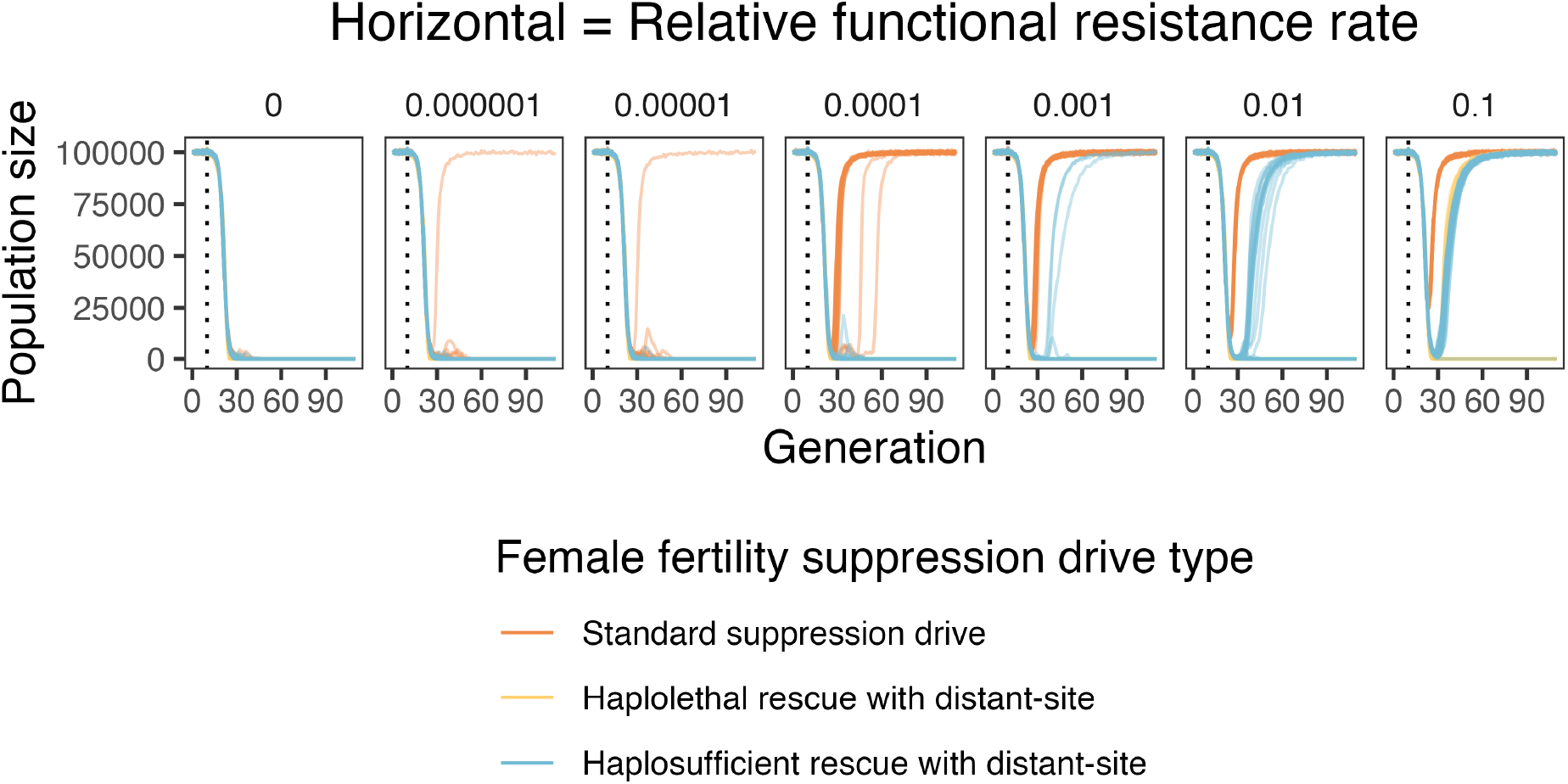
Example population trajectories under varying relative functional resistance rates. Population suppression with varying functional resistance rates. The gene drive is introduced after generation 10 (the dotted line). The introduction frequency of gene drive heterozygotes is 0.01, and the total population size is 100,000. For each combination of parameters, we ran 50 model repetitions for each drive.

**Figure S9.**
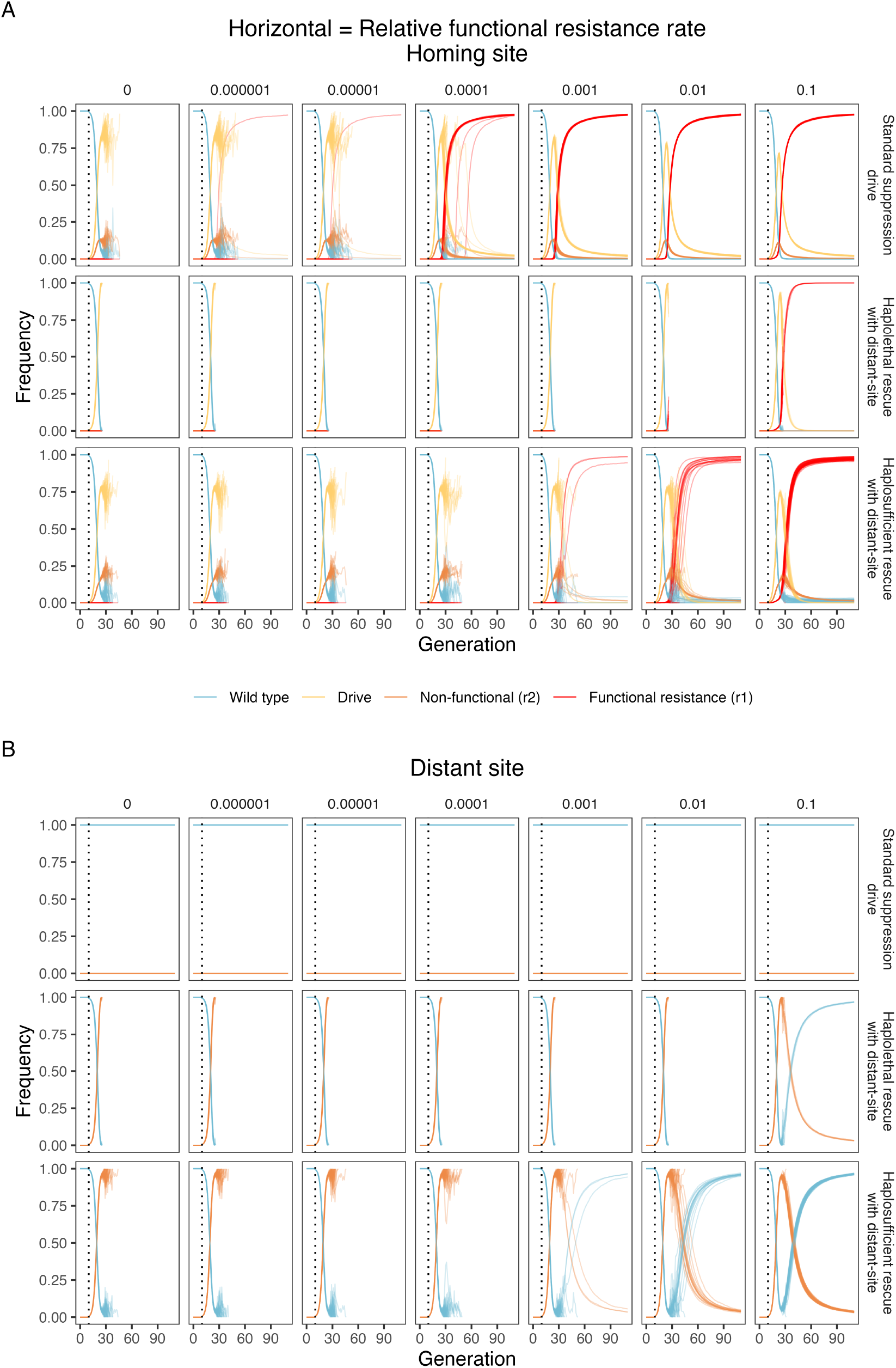
Example allele frequency trajectories. **A)**homing site and **B)** distant site allele frequencies under varying functional resistance rates. The gene drive is introduced after generation 10 (the dotted line). The introduction frequency of gene drive heterozygotes is 0.01, and the total population size is 100,000. For each combination of parameters, we ran 50 model repetitions for each drive.

**Figure S10.**
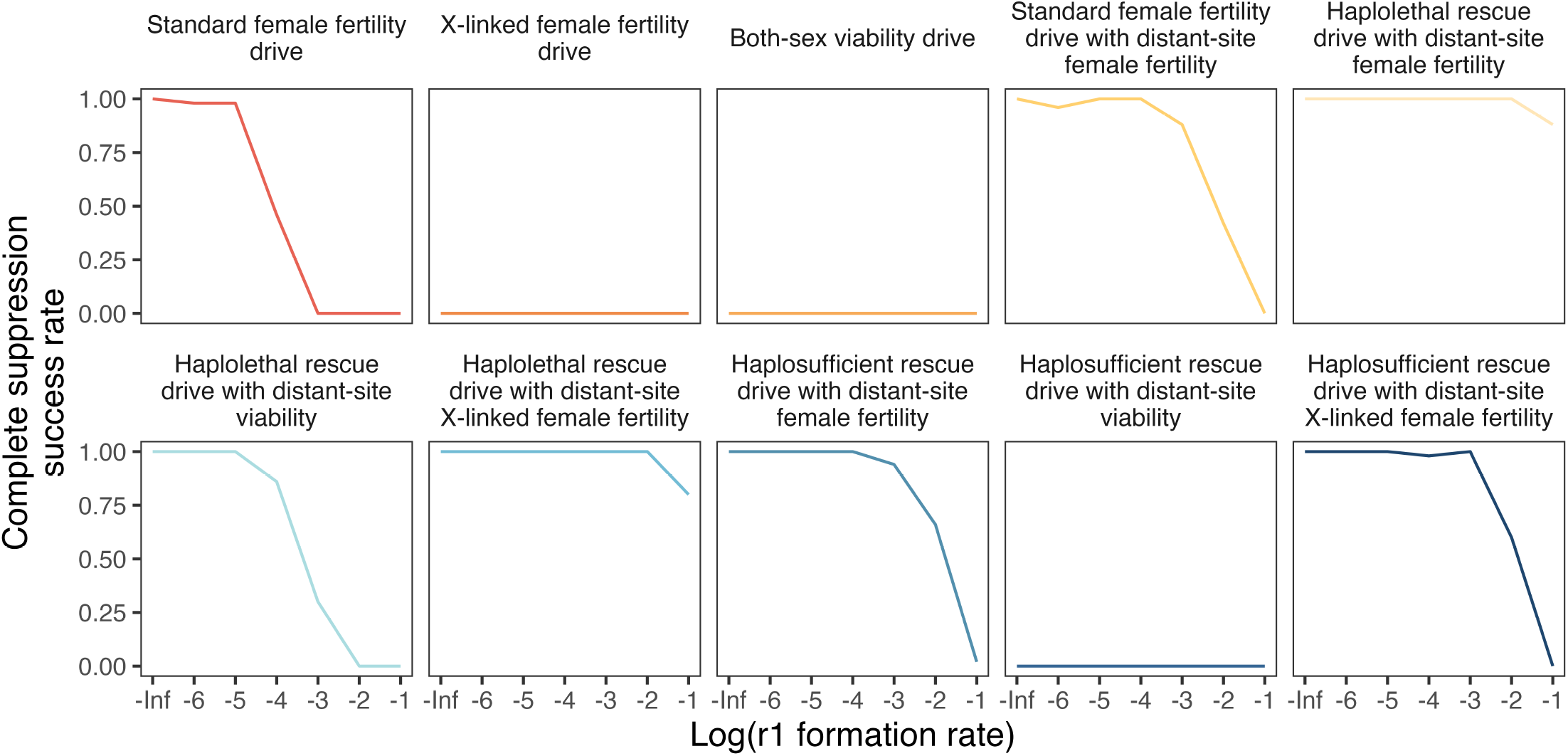
Impact of functional resistance alleles on drive performance. Complete suppression success rate under various functional resistance allele formation rates. The r1 formation rate is the relative rate of resistance alleles that are functional, rather than nonfunctional r2 alleles. The model is run for 100 generations after an introduction of drive heterozygotes at 0.01 frequency into a population of 100,000. We used a total cut rate 1.0, drive conversion rate 0.9, somatic expression female fertility fitness effect of 0.9, and an embryo cut rate of 0.1 in the progeny of female drive carriers. The complete suppression success rate is calculated as the number of repetitions in which complete suppression occurs divided by the total number of repetitions. Because the model stops at 111 generations, a last generation of 111 means that suppression was not successful. For each combination of parameters, we ran 50 model repetitions.

**Figure S11.**
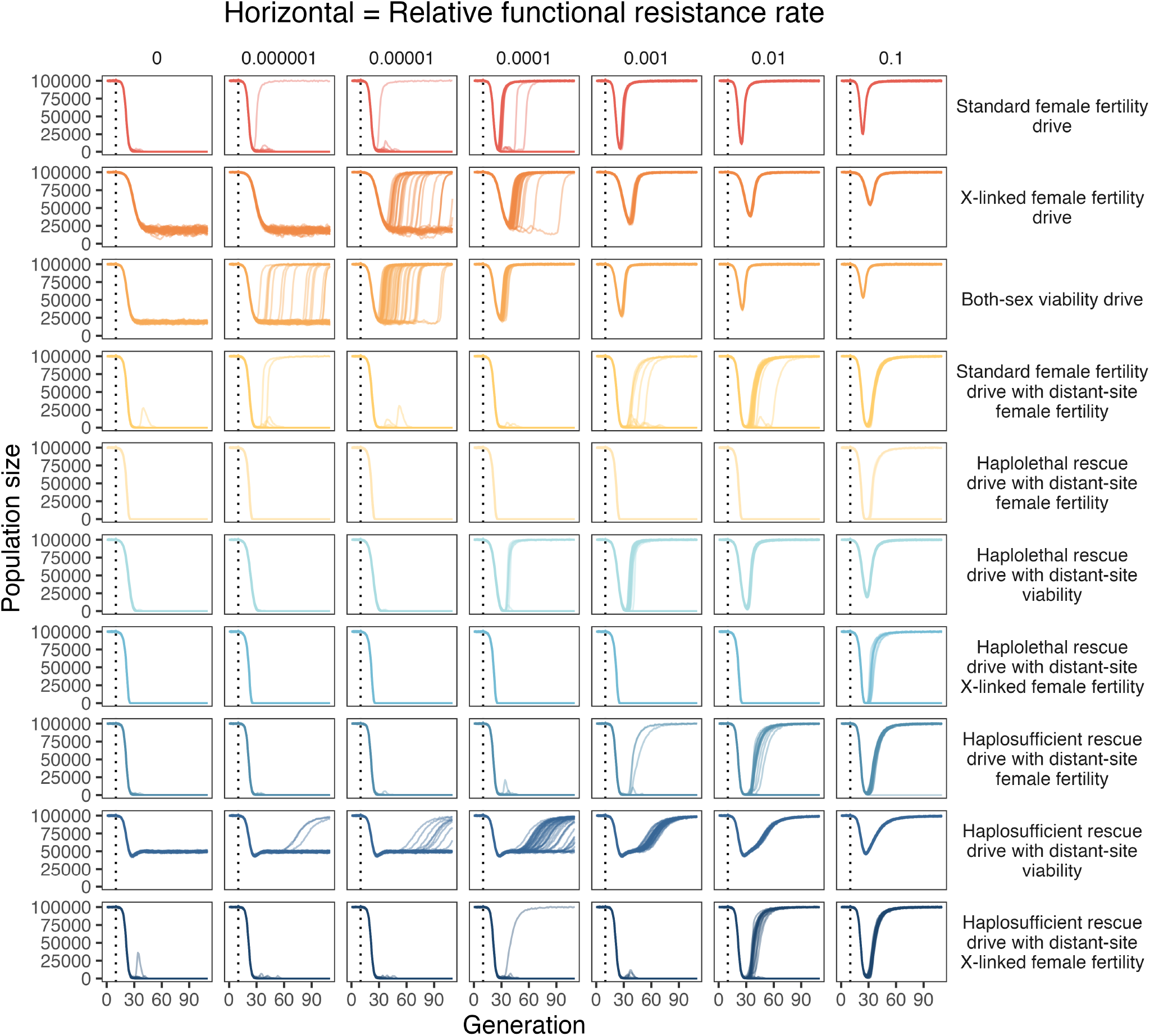
Impact of functional resistance alleles on drive performance. Population suppression with varying functional resistance rates. The r1 formation rate is the relative rate of resistance alleles that are functional, rather than nonfunctional r2 alleles. The model is run for 100 generations after an introduction of drive heterozygotes at 0.01 frequency into a population of 100,000. We used a total cut rate 1.0, drive conversion rate 0.9, somatic expression female fertility fitness effect of 0.9, and an embryo cut rate of 0.1 in the progeny of female drive carriers. For each combination of parameters, we ran 50 model repetitions.

**Figure S12.**
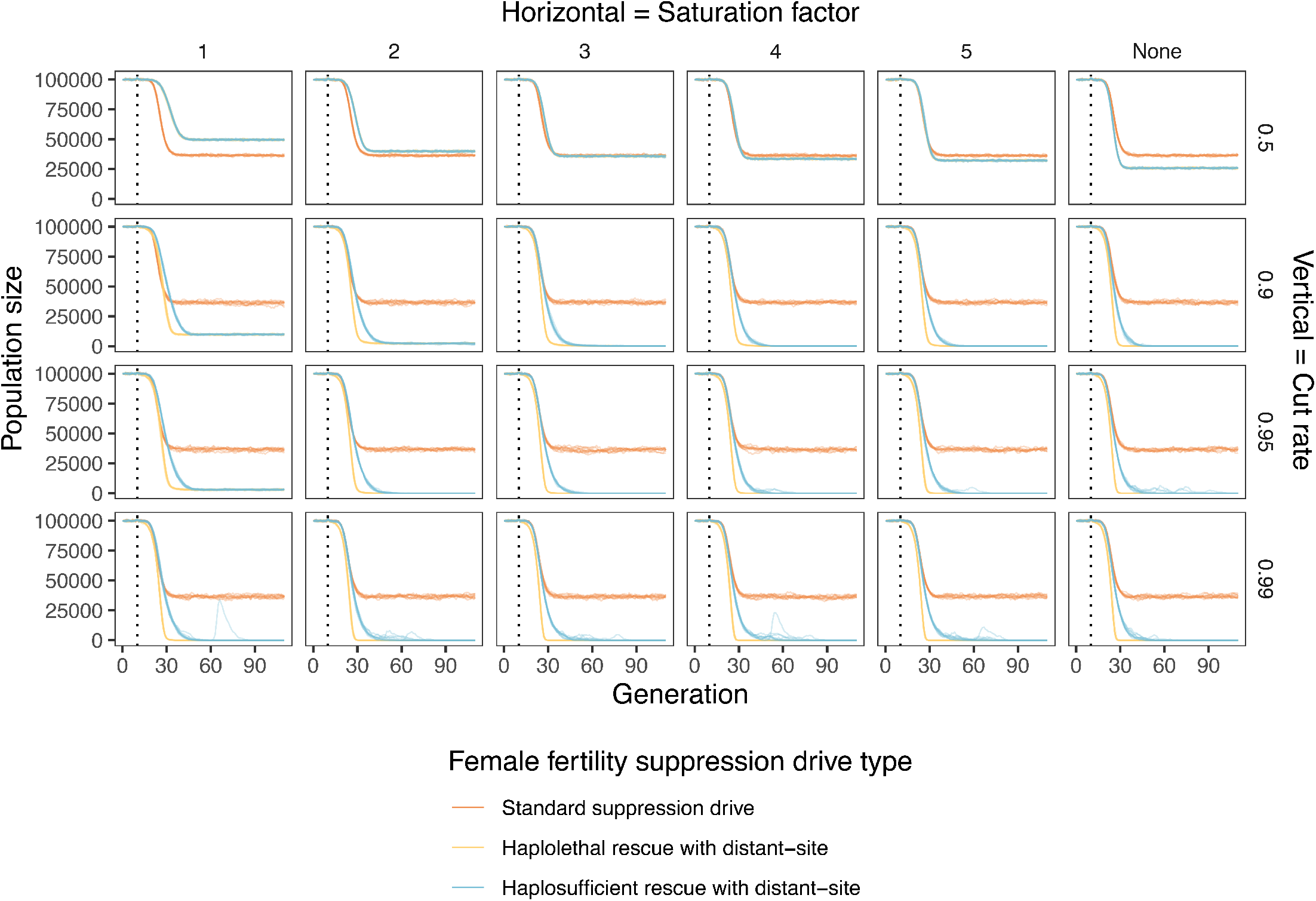
Example population trajectories under varying gRNA saturation factors and total cut rates. The gene drive is introduced after generation 10 (the dotted line). The introduction frequency of gene drive heterozygotes is 0.01, and the total population size is 100,000. We used a drive conversion rate 0.9, somatic expression female fertility fitness effect of 1, and an embryo cut rate of 0 in the progeny of female drive carriers. For each combination of parameters, we ran 10 model repetitions for each drive.

**Figure S13.**
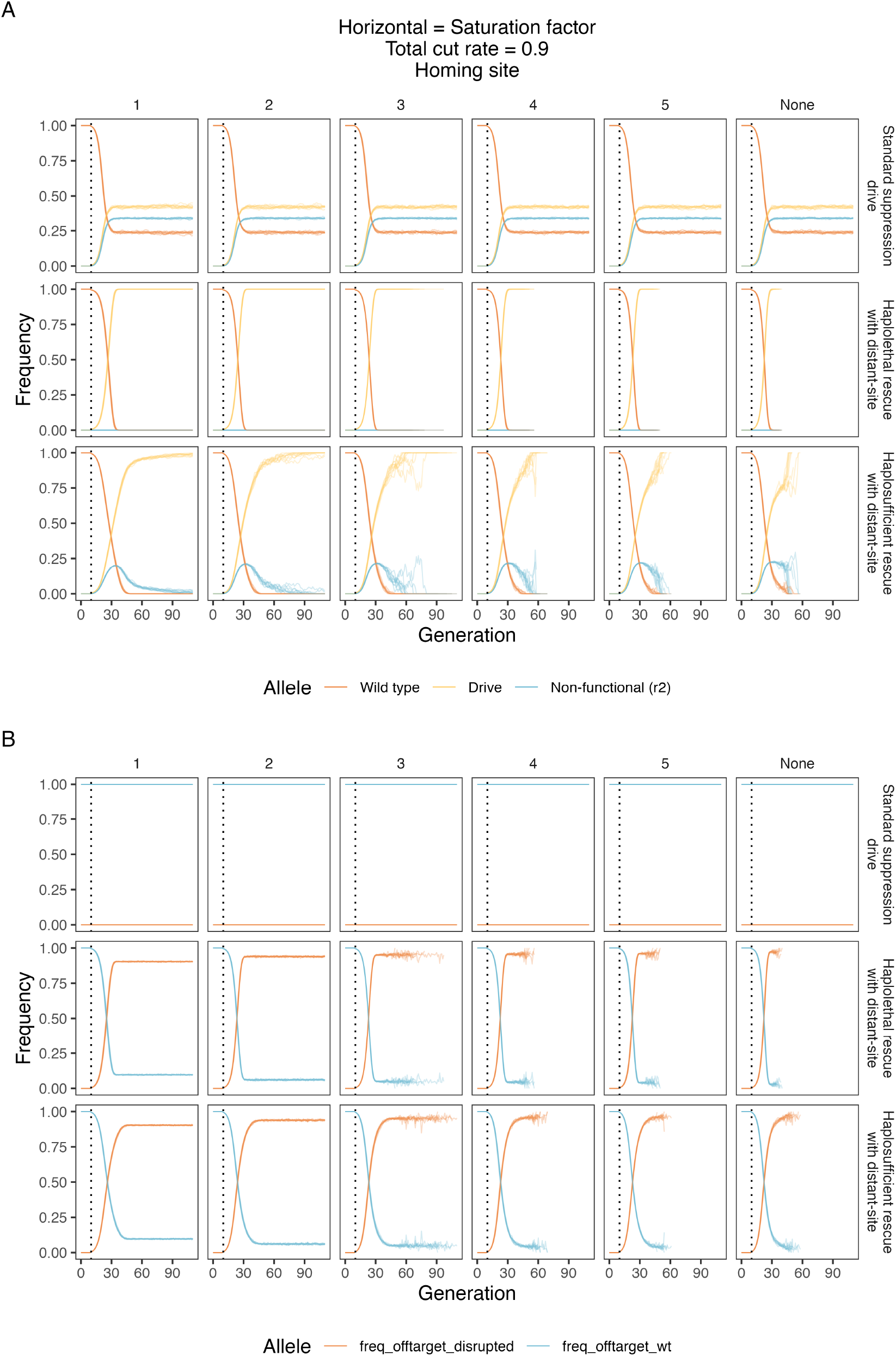
Example allele frequency trajectories. **A)** homing site and **B)** distant site allele frequencies under varying gRNA saturation and total cut rates. The gene drive is introduced after generation 10 (the dotted line). The introduction frequency of gene drive heterozygotes is 0.01, and the total population size is 100,000. We used a drive conversion rate 0.9, somatic expression female fertility fitness effect of 1, and an embryo cut rate of 0 in the progeny of female drive carriers. For each combination of parameters, we ran 10 model repetitions for each drive.

**Figure S14.**
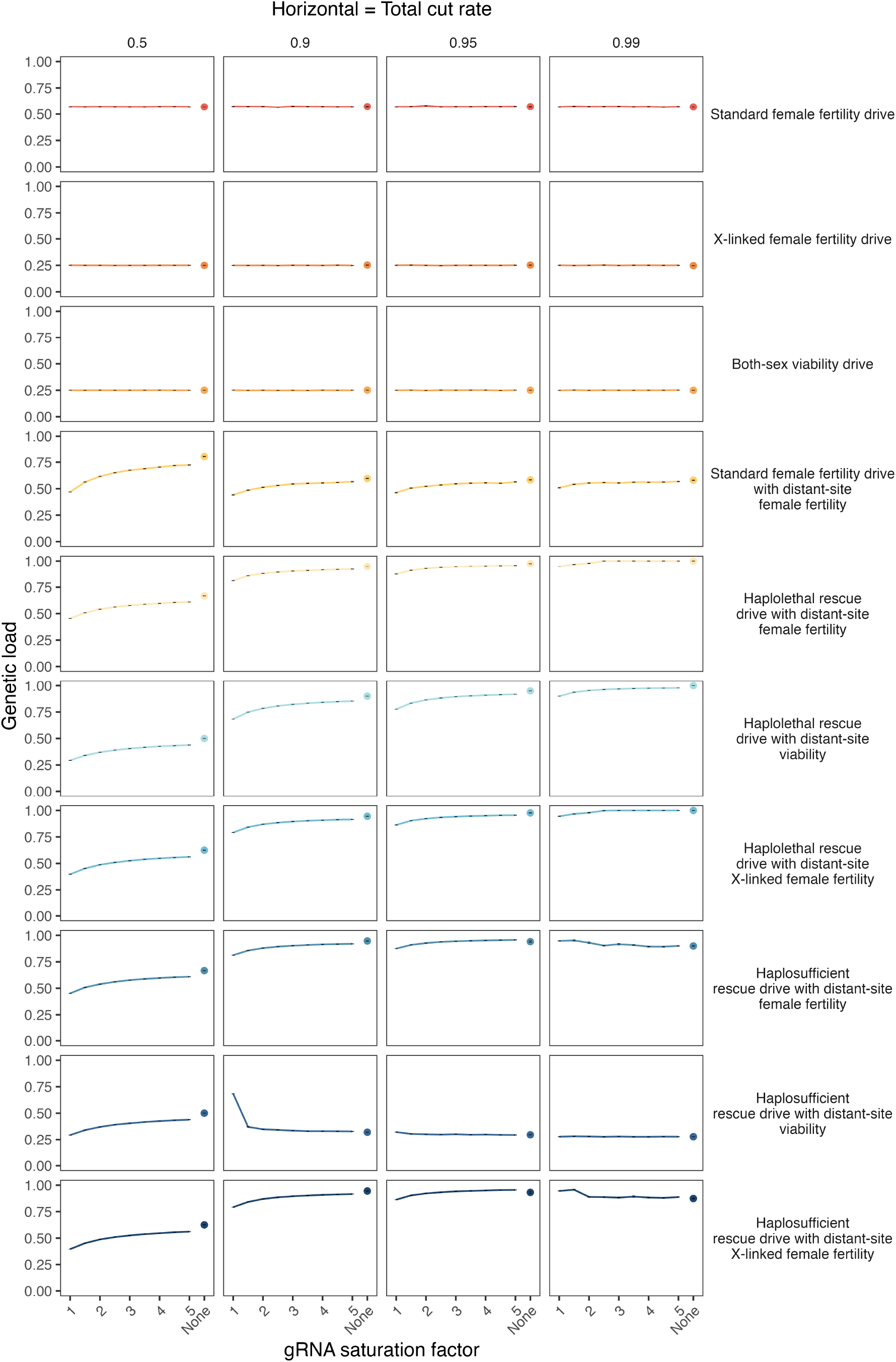
Mean genetic load with variable gRNA saturation factor and total cut rates. The gRNA saturation factor is modelled as relative Cas9 activity with unlimited gRNAs compared to the activity with a single gRNA. We assume that the distant site target gene drives have double the amount of gRNAs compared to a standard suppression drive and that the cut rates are equally reduced at both sites (the two sites are assumed to have the same number of gRNAs). The specified total cut rate is the rate for unlimited gRNA saturation factor. We used a drive conversion rate 0.9, somatic expression female fertility fitness effect of 1, and an embryo cut rate of 0 in the progeny of female drive carriers. For each combination of parameters, we ran 10 model repetitions.

**Figure S15.**
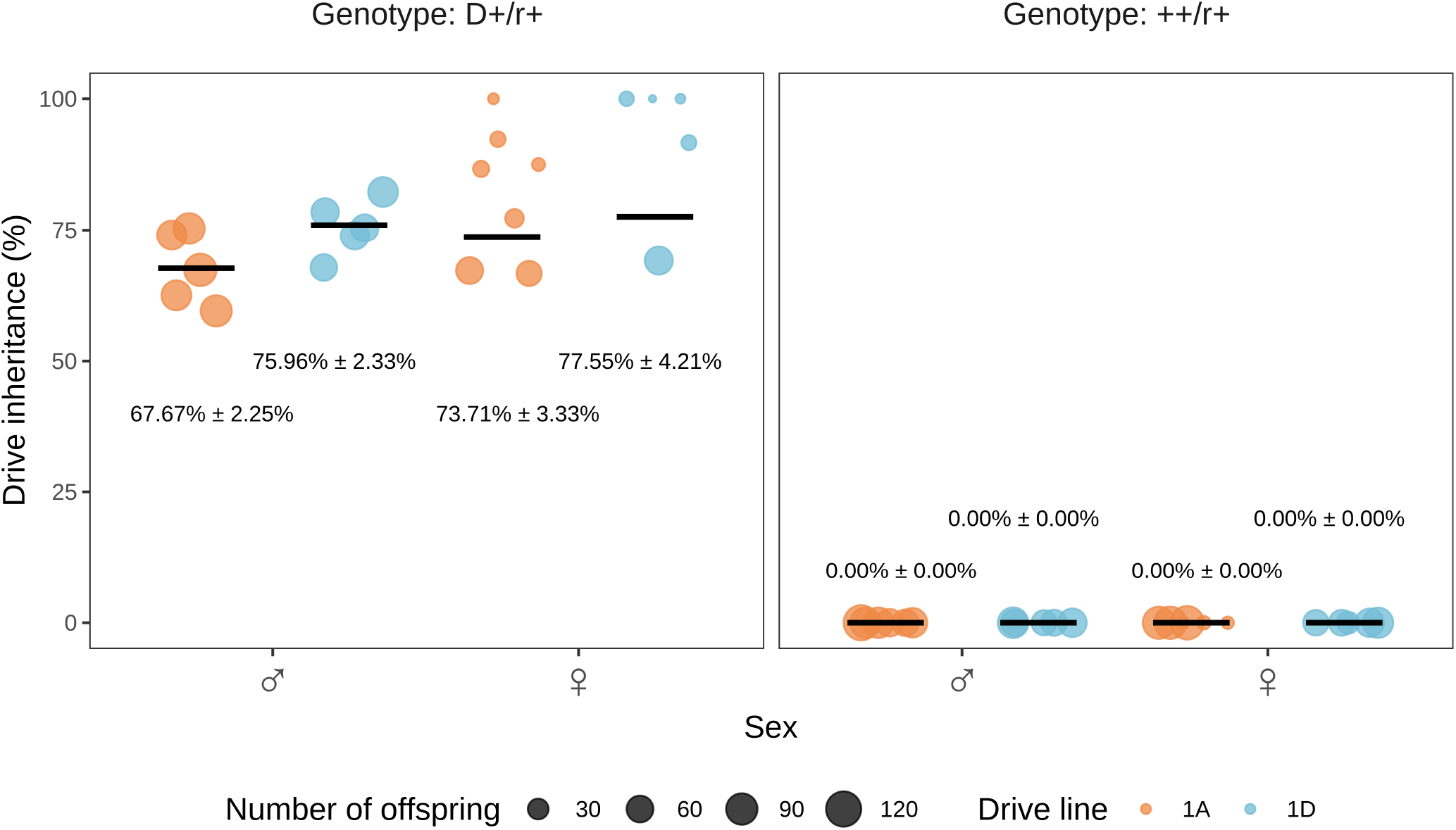
Drive inheritance rates of fertility experiment. The drive inheritance rate indicates the percentage of offspring with EGFP fluorescence from the cross between drive heterozygotes and Cas9 homozygotes with w1118 flies (without drive and Cas9). The size of each dot represents the total number of offspring.

## References

1. Kevin M Esvelt, Andrea L Smidler, Flaminia Catteruccia, and George M Church. Concerning RNA-guided gene drives for the alteration of wild populations. eLife, 3:e03401, July 2014. ISSN 2050-084X. doi: 10.7554/eLife.03401.

2. Jackson Champer, Anna Buchman, and Omar S. Akbari. Cheating evolution: engineering gene drives to manipulate the fate of wild populations. Nature Reviews Genetics, 17(3): 146–159, March 2016. ISSN 1471-0056, 1471-0064. doi: 10.1038/nrg.2015.34.

3. H. Charles J. Godfray, Ace North, and Austin Burt. How driving endonuclease genes can be used to combat pests and disease vectors. BMC Biology, 15(1):81, December 2017. ISSN 1741-7007. doi: 10.1186/s12915-017-0420-4.

4. Ethan Bier. Gene drives gaining speed. Nature Reviews Genetics, 23(1):5–22, January 2022. ISSN 1471-0056, 1471-0064. doi: 10.1038/s41576-021-00386-0.

5. Guan-Hong Wang, Jie Du, Chen Yi Chu, Mukund Madhav, Grant L. Hughes, and Jackson Champer. Symbionts and gene drive: two strategies to combat vector-borne disease. Trends in Genetics, 38(7):708–723, Jul 2022. ISSN 01689525. doi: 10.1016/j.tig.2022.02.013.

6. Luke S. Alphey, Andrea Crisanti, Filippo (Fil) Randazzo, and Omar S. Akbari. Opinion: Standardizing the definition of gene drive. Proceedings of the National Academy of Sciences, page 202020417, November 2020. ISSN 0027-8424, 1091-6490. doi: 10.1073/pnas.2020417117.

7. Valentino M Gantz, Nijole Jasinskiene, Olga Tatarenkova, Aniko Fazekas, Vanessa M Macias, Ethan Bier, and Anthony A James. Highly efficient cas9-mediated gene drive for population modification of the malaria vector mosquito anopheles stephensi. Proceedings of the National Academy of Sciences, 112(49):E6736–E6743, 2015.

8. Kyros Kyrou, Andrew M Hammond, Roberto Galizi, Nace Kranjc, Austin Burt, Andrea K Beaghton, Tony Nolan, and Andrea Crisanti. A CRISPR–Cas9 gene drive targeting doublesex causes complete population suppression in caged Anopheles gambiae mosquitoes. Nature Biotechnology, 36(11):1062–1066, November 2018. ISSN 1087-0156, 1546-1696. doi: 10.1038/nbt.4245.

9. Thomas A. A. Prowse, Phillip Cassey, Joshua V. Ross, Chandran Pfitzner, Talia A. Wittmann, and Paul Thomas. Dodging silver bullets: good CRISPR gene-drive design is critical for eradicating exotic vertebrates. Proceedings of the Royal Society B: Biological Sciences, 284(1860):20170799, August 2017. ISSN 0962-8452, 1471-2954. doi: 10.1098/rspb.2017.0799.

10. Luke Gierus, Aysegul Birand, Mark D. Bunting, Gelshan I. Godahewa, Sandra G. Piltz, Kevin P. Oh, Antoinette J. Piaggio, David W. Threadgill, John Godwin, Owain Edwards, Phillip Cassey, Joshua V. Ross, Thomas A. A. Prowse, and Paul Q. Thomas. Leveraging a natural murine meiotic drive to suppress invasive populations. Proceedings of the National Academy of Sciences, 119(46):e2213308119, November 2022. ISSN 0027-8424, 1091-6490. doi: 10.1073/pnas.2213308119.

11. Kym E. Wilkins, Thomas A.A. Prowse, Phillip Cassey, Paul Q. Thomas, and Joshua V. Ross. Pest demography critically determines the viability of synthetic gene drives for population control. Mathematical Biosciences, 305:160–169, November 2018. ISSN 00255564. doi: 10.1016/j.mbs.2018.09.005.

12. Samuel E. Champer, Nathan Oakes, Ronin Sharma, Pablo García-Díaz, Jackson Champer, and Philipp W. Messer. Modeling CRISPR gene drives for suppression of invasive rodents using a supervised machine learning framework. PLOS Computational Biology, 17(12): e1009660, December 2021. ISSN 1553-7358. doi: 10.1371/journal.pcbi.1009660.

13. Kevin M. Esvelt and Neil J. Gemmell. Conservation demands safe gene drive. PLOS Biology, 15(11):e2003850, November 2017. ISSN 1545-7885. doi: 10.1371/journal.pbio.2003850.

14. Sumit Dhole, Alun L. Lloyd, and Fred Gould. Gene Drive Dynamics in Natural Populations: The Importance of Density Dependence, Space, and Sex. Annual Review of Ecology, Evolution, and Systematics, 51(1):505–531, November 2020. ISSN 1543-592X, 1545-2069. doi: 10.1146/annurev-ecolsys-031120-101013.

15. Tom A R Price, Nikolai Windbichler, Robert L Unckless, Andreas Sutter, Jan-Niklas Runge, Perran A Ross, Andrew Pomiankowski, Nicole L Nuckolls, Catherine Montchamp-Moreau, Nicole Mideo, Oliver Y Martin, Andri Manser, Mathieu Legros, Amanda M Larracuente, Luke Holman, John Godwin, Neil Gemmell, Cécile Courret, Anna Buchman, Luke G Barrett, and Anna K Lindholm. Resistance to natural and synthetic gene drive systems. Journal of evolutionary biology, 33(10):1345–1360, 2020.

16. Andrew M. Hammond, Kyros Kyrou, Marco Bruttini, Ace North, Roberto Galizi, Xenia Karlsson, Nace Kranjc, Francesco M. Carpi, Romina D’Aurizio, Andrea Crisanti, and Tony Nolan. The creation and selection of mutations resistant to a gene drive over multiple generations in the malaria mosquito. PLOS Genetics, 13(10):e1007039, October 2017. ISSN 1553-7404. doi: 10.1371/journal.pgen.1007039.

17. Chandran Pfitzner, Melissa A. White, Sandra G. Piltz, Michaela Scherer, Fatwa Adikusuma, James N. Hughes, and Paul Q. Thomas. Progress Toward Zygotic and Germline Gene Drives in Mice. The CRISPR Journal, 3(5):388–397, October 2020. ISSN 2573-1599, 2573-1602. doi: 10.1089/crispr.2020.0050.

18. Hannah A. Grunwald, Valentino M. Gantz, Gunnar Poplawski, Xiang-Ru S. Xu, Ethan Bier, and Kimberly L. Cooper. Super-Mendelian inheritance mediated by CRISPR–Cas9 in the female mouse germline. Nature, 566(7742):105–109, February 2019. ISSN 0028-0836, 1476-4687. doi: 10.1038/s41586-019-0875-2.

19. Samuel E. Champer, Suh Yeon Oh, Chen Liu, Zhaoxin Wen, Andrew G. Clark, Philipp W. Messer, and Jackson Champer. Computational and experimental performance of CRISPR homing gene drive strategies with multiplexed gRNAs. Science Advances, 6(10):eaaz0525, March 2020. ISSN 2375-2548. doi: 10.1126/sciadv.aaz0525.

20. Jackson Champer, Riona Reeves, Suh Yeon Oh, Chen Liu, Jingxian Liu, Andrew G. Clark, and Philipp W. Messer. Novel CRISPR/Cas9 gene drive constructs reveal insights into mechanisms of resistance allele formation and drive efficiency in genetically diverse populations. PLOS Genetics, 13(7):e1006796, July 2017. ISSN 1553-7404. doi: 10.1371/journal.pgen.1006796.

21. Eli M. Carrami, Kolja N. Eckermann, Hassan M. M. Ahmed, Héctor M. Sánchez C., Stefan Dippel, John M. Marshall, and Ernst A. Wimmer. Consequences of resistance evolution in a cas9-based sex conversion-suppression gene drive for insect pest management. Proceedings of the National Academy of Sciences, 115(24):6189–6194, Jun 2018. ISSN 0027-8424, 1091-6490. doi: 10.1073/pnas.1713825115.

22. Thai Binh Pham, Celine Hien Phong, Jared B. Bennett, Kristy Hwang, Nijole Jasinskiene, Kiona Parker, Drusilla Stillinger, John M. Marshall, Rebeca Carballar-Lejarazú, and Anthony A. James. Experimental population modification of the malaria vector mosquito, anopheles stephensi. PLOS Genetics, 15(12):e1008440, Dec 2019. ISSN 1553-7404. doi: 10.1371/journal.pgen.1008440.

23. Nicolas O. Rode, Arnaud Estoup, Denis Bourguet, Virginie Courtier-Orgogozo, and Florence Débarre. Population management using gene drive: molecular design, models of spread dynamics and assessment of ecological risks. Conservation Genetics, 20(4):671–690, Aug 2019. ISSN 1566-0621, 1572-9737. doi: 10.1007/s10592-019-01165-5.

24. Georg Oberhofer, Tobin Ivy, and Bruce A Hay. Behavior of homing endonuclease gene drives targeting genes required for viability or female fertility with multiplexed guide rnas. Proceedings of the National Academy of Sciences, 115(40):E9343–E9352, 2018.

25. Emily Yang, Matthew Metzloff, Anna M Langmüller, Xuejiao Xu, Andrew G Clark, Philipp W Messer, and Jackson Champer. A homing suppression gene drive with multiplexed grnas maintains high drive conversion efficiency and avoids functional resistance alleles. G3, 12 (6):jkac081, 2022.

26. Andrew Hammond, Xenia Karlsson, Ioanna Morianou, Kyros Kyrou, Andrea Beaghton, Matthew Gribble, Nace Kranjc, Roberto Galizi, Austin Burt, Andrea Crisanti, and Tony Nolan. Regulating the expression of gene drives is key to increasing their invasive potential and the mitigation of resistance. PLOS Genetics, 17(1):e1009321, Jan 2021. ISSN 1553-7404. doi: 10.1371/journal.pgen.1009321.

27. Rebeca Carballar-Lejarazú, Christian Ogaugwu, Taylor Tushar, Adam Kelsey, Thai Binh Pham, Jazmin Murphy, Hanno Schmidt, Yoosook Lee, Gregory C. Lanzaro, and Anthony A. James. Next-generation gene drive for population modification of the malaria vector mosquito, anopheles gambiae. Proceedings of the National Academy of Sciences, 117(37): 22805–22814, Sep 2020. ISSN 0027-8424, 1091-6490. doi: 10.1073/pnas.2010214117.

28. Jie Du, Weizhe Chen, Xihua Jia, Xuejiao Xu, Emily Yang, Ruizhi Zhou, Yuqi Zhang, Matt Metzloff, Philipp W. Messer, and Jackson Champer. New germline cas9 promoters show improved performance for homing gene drive. BioRxiv, Jul 2023. doi: 10.1101/2023.07.16.549205.

29. Andrew Hammond, Roberto Galizi, Kyros Kyrou, Alekos Simoni, Carla Siniscalchi, Dimitris Katsanos, Matthew Gribble, Dean Baker, Eric Marois, Steven Russell, Austin Burt, Nikolai Windbichler, Andrea Crisanti, and Tony Nolan. A CRISPR-Cas9 gene drive system targeting female reproduction in the malaria mosquito vector Anopheles gambiae. Nature Biotechnology, 34(1):78–83, January 2016. ISSN 1087-0156, 1546-1696. doi: 10.1038/nbt.3439.

30. Ming Li, Ting Yang, Nikolay P Kandul, Michelle Bui, Stephanie Gamez, Robyn Raban, Jared Bennett, Héctor M Sánchez C, Gregory C Lanzaro, Hanno Schmidt, Yoosook Lee, John M Marshall, and Omar S Akbari. Development of a confinable gene drive system in the human disease vector aedes aegypti. eLife, 9:e51701, Jan 2020. ISSN 2050-084X. doi: 10.7554/eLife.51701.

31. William Reid, Adeline E Williams, Irma Sanchez-Vargas, Jingyi Lin, Rucsanda Juncu, Ken E Olson, and Alexander W E Franz. Assessing single-locus crispr/cas9-based gene drive variants in the mosquito aedes aegypti via single-generation crosses and modeling. G3 Genes|Genomes|Genetics, 12(12):jkac280, Dec 2022. ISSN 2160-1836. doi: 10.1093/g3journal/jkac280.

32. Michelle A. E. Anderson, Estela Gonzalez, Joshua X. D. Ang, Lewis Shackleford, Katherine Nevard, Sebald A. N. Verkuijl, Matthew P. Edgington, Tim Harvey-Samuel, and Luke Alphey. Closing the gap to effective gene drive in aedes aegypti by exploiting germline regulatory elements. Nature Communications, 14(1):338, Jan 2023. ISSN 2041-1723. doi: 10.1038/s41467-023-36029-7.

33. Yuk-Sang Chan, David S. Huen, Ruth Glauert, Eleanor Whiteway, and Steven Russell. Optimising homing endonuclease gene drive performance in a semi-refractory species: The drosophila melanogaster experience. PLoS ONE, 8(1):e54130, Jan 2013. ISSN 1932-6203. doi: 10.1371/journal.pone.0054130.

34. Rebeca Carballar-Lejarazú, Taylor Tushar, Thai Binh Pham, and Anthony A James. Cas9-mediated maternal effect and derived resistance alleles in a gene-drive strain of the african malaria vector mosquito, anopheles gambiae. Genetics, page iyac055, Apr 2022. ISSN 1943-2631. doi: 10.1093/genetics/iyac055.

35. Amarish K. Yadav, Cole Butler, Akihiko Yamamoto, Anandrao A. Patil, Alun L. Lloyd, and Maxwell J. Scott. Crispr/cas9-based split homing gene drive targeting doublesex for population suppression of the global fruit pest drosophila suzukii. Proceedings of the National Academy of Sciences, 120(25):e2301525120, Jun 2023. ISSN 0027-8424, 1091-6490. doi: 10.1073/pnas.2301525120.

36. Jackson Champer, Emily Yang, Esther Lee, Jingxian Liu, Andrew G. Clark, and Philipp W. Messer. A crispr homing gene drive targeting a haplolethal gene removes resistance alleles and successfully spreads through a cage population. Proceedings of the National Academy of Sciences, 117(39):24377–24383, Sep 2020. ISSN 0027-8424, 1091-6490. doi: 10.1073/pnas.2004373117.

37. Nikolay P Kandul, Junru Liu, Anna Buchman, Valentino M Gantz, Ethan Bier, and Omar S Akbari. Assessment of a Split Homing Based Gene Drive for Efficient Knockout of Multiple Genes. G3 Genes|Genomes|Genetics, 10(2):827–837, 02 2020. ISSN 2160-1836. doi: 10.1534/g3.119.400985.

38. Benjamin C. Haller and Philipp W. Messer. SLiM 4: Multispecies eco-evolutionary modeling. The American Naturalist, page 723601, November 2022. ISSN 0003-0147, 1537-5323. doi: 10.1086/723601.

39. R Core Team. R: A Language and Environment for Statistical Computing. R Foundation for Statistical Computing, Vienna, Austria, 2021.

40. M Wayne Davis and Erik M Jorgensen. Ape, a plasmid editor: a freely available dna manipulation and visualization program. Frontiers in Bioinformatics, 2:818619, 2022.

41. Jackson Champer, Samuel E. Champer, Isabel K. Kim, Andrew G. Clark, and Philipp W. Messer. Design and analysis of crispr-based underdominance toxin-antidote gene drives. Evolutionary Applications, 14(4):1052–1069, Apr 2021. ISSN 1752-4571, 1752-4571. doi: 10.1111/eva.13180.

42. Jackson Champer, Isabel K. Kim, Samuel E. Champer, Andrew G. Clark, and Philipp W. Messer. Performance analysis of novel toxin-antidote crispr gene drive systems. BMC Biology, 18(1):27, Dec 2020. ISSN 1741-7007. doi: 10.1186/s12915-020-0761-2.

43. Siân AM Spinner, Zoe H Barnes, Alin Mirel Puinean, Pam Gray, Tarig Dafa’alla, Caroline E Phillips, Camila Nascimento de Souza, Tamires Fonseca Frazon, Kyla Ercit, Amandine Collado, et al. New self-sexing aedes aegypti strain eliminates barriers to scalable and sustainable vector control for governments and communities in dengue-prone environments. Frontiers in Bioengineering and Biotechnology, 10, 2022.

44. Yiran Liu and Jackson Champer. Modelling homing suppression gene drive in haplodiploid organisms. Proceedings of the Royal Society B, 289(1972):20220320, 2022.

45. Pavel P Khil, Natalya A Smirnova, Peter J Romanienko, and R Daniel Camerini-Otero. The mouse x chromosome is enriched for sex-biased genes not subject to selection by meiotic sex chromosome inactivation. Nature Genetics, 36(6):642–646, Jun 2004. ISSN 1061-4036, 1546-1718. doi: 10.1038/ng1368.

46. Michael Parisi, Rachel Nuttall, Daniel Naiman, Gerard Bouffard, James Malley, Justen Andrews, Scott Eastman, and Brian Oliver. Paucity of genes on the drosophila x chromosome showing male-biased expression. Science, 299(5607):697–700, Jan 2003. ISSN 0036-8075, 1095-9203. doi: 10.1126/science.1079190.

